# A universal subcuticular bacterial symbiont of a coral predator, the crown-of-thorns starfish, in the Indo-Pacific

**DOI:** 10.1101/2020.02.13.947093

**Authors:** Naohisa Wada, Hideaki Yuasa, Rei Kajitani, Yasuhiro Gotoh, Yoshitoshi Ogura, Dai Yoshimura, Atsushi Toyoda, Sen-Lin Tang, Yukio Higashimura, Hugh Sweatman, Zac Forsman, Omri Bronstein, Gal Eyal, Nalinee Thongtham, Takehiko Itoh, Tetsuya Hayashi, Nina Yasuda

## Abstract

**Background:** Population outbreaks of the crown-of-thorns starfish (*Acanthaster planci* sensu lato; COTS), a primary predator of reef-building corals in the Indo-Pacific Ocean, are major concerns in coral reef management. While biological and ecological knowledge of COTS has been accumulating since the 1960s, little is known about its associated bacteria. The aim of this study was to provide fundamental information on dominant COTS-associated bacteria through a multifaceted molecular approach.

**Methods:** A total of 205 COTS individuals from 17 locations throughout the Indo-Pacific Ocean were examined for the presence of COTS-associated bacteria. We conducted 16S rRNA metabarcoding of COTS to determine the bacterial profiles of different parts of the body, and generated a full-length 16S rRNA gene sequence from a single dominant bacterium, which we designated COTS27. We performed phylogenetic analysis to determine the taxonomy, screening of COTS27 across the Indo-Pacific, FISH to visualize it within the COTS tissues, and reconstruction of the chromosome from the hologenome sequence data.

**Results:** We discovered that a single bacterium exists at high densities in the subcuticular space in COTS forming a biofilm-like structure between the cuticle and the epidermis. COTS27 belongs to a clade that presumably represents a distinct order (so-called marine spirochetes) in the phylum *Spirochaetes* and is universally present in COTS throughout the Indo-Pacific Ocean. The reconstructed genome of COTS27 includes some genetic traits that are probably linked to adaptation to marine environments and evolution as an extracellular endosymbiont in subcuticular spaces.

**Conclusions:** COTS27 can be found in three allopatrically speciated COTS species, ranging from northern Red Sea to the Pacific, implying that symbiotic relationship arose before the speciation (approximately 2 million years ago). The universal association of COTS27 with COTS and nearly mono-specific association at least with the Indo-Pacific COTS potentially provides a useful model system for studying symbiont-host interactions in marine invertebrates.

## Introduction

Coral reefs support almost one-third of the world’s marine coastal species [1,2]. However, population outbreaks of a coral predator, the crown-of-thorns starfish (*Acanthaster planci* sensu lato; COTS), are a great threat to Indo-Pacific coral reef ecosystem integrity and biodiversity [3–5]. A 27-year study of the Great Barrier Reef concluded that COTS outbreaks and tropical cyclones were the main causes of the loss of reef-building corals (1985-2012) [6]. While some aspects of the biology of COTS, such as its reproduction, larval ecology, phylogeography, and behaviour, have been studied intensively [5], little is known about its associated microbiota.

The bacterial symbionts of marine invertebrates have been shown to be important to their host organisms [7]. In echinoderms, bacterial communities may play a role in larval settlement [8], amino acid uptake on the integument [9], and digestive strategies in the gut [10,11], and these communities may even drive morphological variations in their host [12]. Bacterial symbionts are prevalent on the body surfaces of echinoderms [13], showing high host specificity [14,15]. Notably, extracellular endosymbionts known as subcuticular bacteria (SCB [16]) have been shown to reside under the cuticular layer of echinoderm fauna from all five extant classes, and it has been postulated that these bacteria provide dissolved free amino acids to their echinoderm hosts [9,17]. To date, molecular genetic approaches targeting the 16S rRNA gene have revealed that several proteobacteria (*Alphaproteobacteria* and *Gammaproteobacteria*) are SCB that are distributed in the subcuticular space in two brittle star species [13,18], one holothurian species [19], and one asteroid species (19).

Despite their potential biological importance, the studies of the bacteria associated with COTS have been mostly culture-based, and only two culture-independent studies have been published to date. Carrier et al. reported shifts in the COTS larval microbiomes associated with diet [20]. Høj et al. found that adult COTS exhibit tissue-specific bacterial communities, largely comprising four major bacterial groups: *Mollicutes* in male gonads*, Spirochaetales* in the body wall*, Hyphomonadaxeae* in the tube feet, and *Oceanospirillales* in all tissues [21]. Although these studies significantly increased our understanding of the COTS microbiome, there is still a great lack of knowledge regarding COTS-associated bacteria, particularly SCB, despite being common in many echinoderm taxa, where they may play an important role for their host organisms.

In the current study, we aimed to obtain primary information on the indigenous bacteria of the body surface of COTS. We carried out a comprehensive analysis of bacterial symbionts associated with COTS in a total of 205 individuals collected from the northern Red Sea to the Pacific over a 13-years period. We highlighted the existence of dominant SCB in COTS, its novel phylogenetic status, universal distribution in the Indo-Pacific COTS, and its genomic characteristics, all of which provide insights into interactions between the COTS host and the SCB.

## Results

### Identification of a single OTU (COTS27) that dominates the body surface microbiota of COTS using 16S rRNA metabarcoding analysis

We used 16S rRNA metabarcoding to analyse the bacterial composition of the microbiota in the body parts (7-8 body parts; disc spines [top and base], arm spines [top and base], ambulacral spines [top and base for Okinawa, or the whole spine for Miyazaki], tube feet, and pyloric stomachs; **Fig.1b**) of six COTS individuals that were collected in Miyazaki and Okinawa, Japan (three individuals from each location). Seawater samples from the same locations were similarly analysed for their bacterial compositions (three samples from each location). After quality filtering, 1,427,570 sequences of bacterial origins were obtained from the COTS samples (n=130 for all body parts in replicates or duplicates; **Suppl. table S1**) and 108,334 bacterial sequences from seawater samples (n=6) with an averages of 10,981 and 18,056 sequences per sample, respectively (**Suppl. table S2**). From the abovementioned sequences, 671 bacterial OTUs were identified, 503 and 401 of which were found in the COTS and seawater samples, respectively. There were 233 OTUs that were common to both. The OTUs that were identified in the COTS and seawater samples represented the bacterial taxa with 144 and 96 families, 29 and 22 OTUs were unclassified, 7 and 12 OTUs were as unknown (classified as bacteria by Silva SINA [22]), respectively (see more detail in **Suppl. table S3**). The rarefaction curves based on the OTUs indicated that all samples reached saturation points (**Suppl. fig. S1**).

**Fig. 1.**
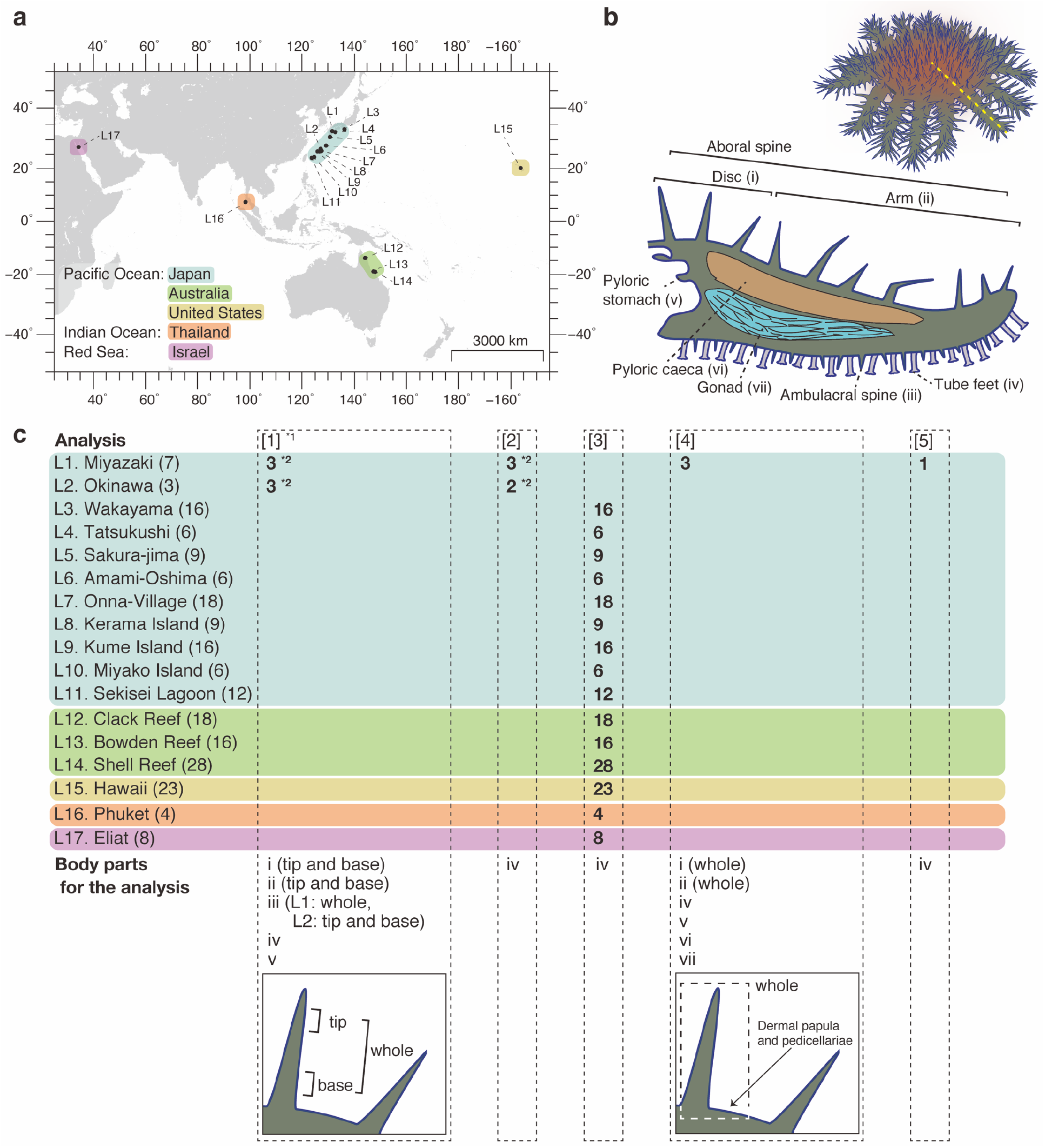
Geographic and anatomical distributions of COTS individuals and the COTS body parts analysed in this study. The seventeen locations where the COTS individuals were collected **(a**) and the dissected body parts of COTS for the analyses (**b**) are shown. The dashed yellow line (panel **b**) indicates the dissection line for the cross-sectional view. In panel (**c**), details of the samples used in each analysis are shown: [1] 16S rRNA metabarcoding, [2] phylogenetic analysis using the full-length 16S rRNA gene sequences, [3] PCR screening and sequencing of the 16S rRNA gene sequences of COTS27, [4] FISH analysis, and [5] hologenome sequencing analysis. *1: This analysis was performed in triplicate for each sample. *2: The same individuals were used in analyses [1] and [2].

In the six COTS individuals that were examined, the relative abundance showed that a single unclassified OTU (OTU 1) occupied 61.8% of the total sequences on average, predominantly in most body parts of both the Okinawan and Miyazaki COTS populations (60.3% and 63.8% of the total sequences on average were assigned to OTU 1 in the Okinawa and Miyazaki COTS collections, respectively; **Fig. 2**), despite the fact that these populations were separated by more than 720 km separating these populations. The high abundance of OTU 1 in all individuals was attributed to the surface body parts (68.8% and 79.1% of the sequences from all spine and tube foot samples, respectively), with 8.0% of these sequences originating from the pyloric stomach samples (**Fig. 2** and **Suppl. table S4**). OTU 1 was abundant at both the aboral (discs and arm spines) and oral (ambulacral spines and tube feet) sides (**Suppl. fig. S2** and **Suppl. table S4**) of the COTS. The tips and bases of the spines showed roughly the same levels of OTU 1 abundance (**Suppl. fig. S2** and **Suppl. table S4**). Five of the 88 spine samples that were examined (containing both tip and base) exhibited no or only a low abundance of OTU 1 (**Suppl. fig. S2**); however, OTU 1 was abundant in the other two DNA preparations of the triplicates from the same sample in all cases, suggesting that the exceptional data from the five preparations were due to some technical problems. OTU 1 was only detected in the Okinawan seawater samples, in which it showed a low abundance (0.026%; **Suppl. fig. S2**). The relatively abundant bacteria other than OTU 1 are described in **Appendix 1**. In total, we identified 41 different OTUs, including OTU 1, in all COTS individuals from the two locations, and these OTUs may represent the core members of the bacterial community of COTS (**Suppl. fig. S3**). The core bacterial OTUs other than OTU 1 accounted for up to 18.4% (the abundance of each OTU was less than 3.5%) of the total reads from all COTS samples (**Suppl. fig. S3d**). These results indicate that a single bacterium (OTU 1) predominantly colonizes the body surface of COTS.

**Fig. 2.**
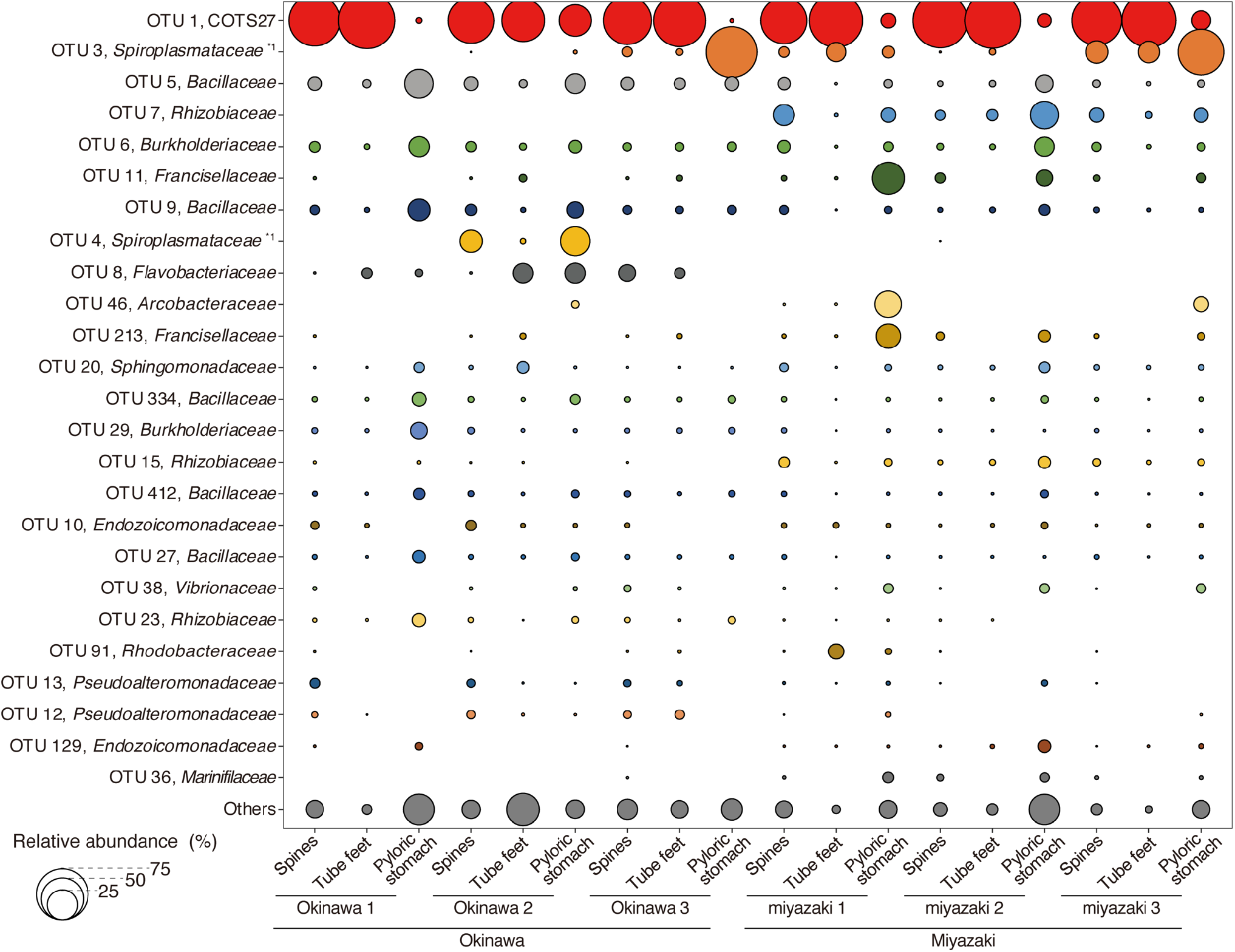
The relative abundances of the 25 most abundant OTUs, including COTS27 (OTU 1; red), in the total samples analysed in this study. The bubble chart of the relative abundances was calculated from the merged replicates of each body part (spines, tube feet, and pyloric stomachs) in each COTS individual. The phylogenies of each OTU were determined based on the results (best hit) of BLAST searches against the NCBI nr/nt database. *1: The phylogenies of OTU3 and OTU4 were determined in the All-species Living Tree Project and RDP databases, respectively.

### Phylogeny of the dominant OTU 1 (COTS27) based on 16S rRNA gene sequences

To elucidate the phylogenetic status of the dominant OTU 1, we determined the full-length 16S rRNA gene sequences of OTU 1 in five tube foot samples obtained from Miyazaki (n=3) and Okinawa (n=2). The five sequences were largely identical (99.9-100% similarity), and there was a partial sequence overlap with the 16S rRNA gene sequence of a spirochete-like bacterium (GenBank accession No. PRJNA420398) that was a dominant bacterium on the body wall of COTS from the Great Barrier Reef [21]. The maximum likelihood (ML) phylogenetic tree (**Fig. 3a**) based on full-length 16S rRNA gene sequences showed that the five sequences related the OTU 1 formed a distinct subclade within one of the three clades of the unclassified spirochete cluster (named clade I; **Fig. 3a**). All sequences in this unclassified spirochete cluster originated from marine environments and marine invertebrates (see **Appendix 2** for more details of clade I) with the exception of a single sequence obtained from a wetland soil sample (GenBank accession No. FQ660021.1). Hereafter, we refer to these spirochetes as “marine spirochetes”, as referred to by Høj et al. (2018) [21]. These marine spirochetes formed a distinct cluster within the phylum *Spirochaetes*, with the order *Brachyspirales* being their closest relative (**Fig. 3a**). Notably, the 16S rRNA gene sequences of the marine spirochetes, including the OTU 1 group, showed only a 76.3–78.1% identity to those of the order *Brachyspirales*, which is well below the proposed threshold for defining a novel order (82.0%) [23]. Thus, the marine spirochetes most likely represent a distinct order in the phylum *Spirochaetes*. Hereafter, we refer to the bacterium corresponding to the OTU 1 as COTS27.

**Fig. 3.**
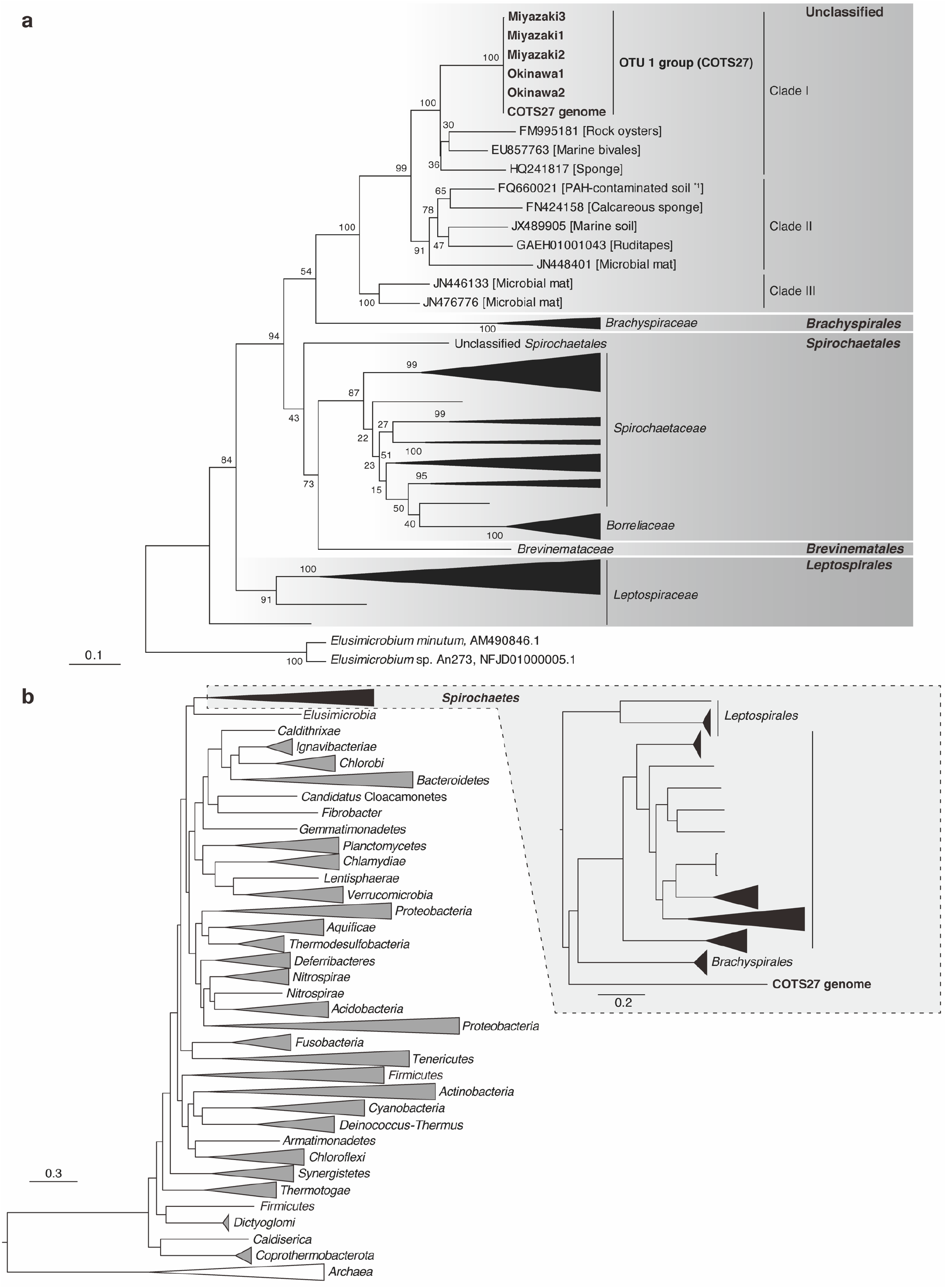
The phylogenetic position of the dominant OTU 1 (COTS27) in the phylum *Spirochaetes*. Maximum likelihood (ML) trees were constructed based on the full-length 16S rRNA gene sequences (**a**) and the sequences of 43 conserved marker genes identified by CheckM (**b**). The bootstrap values in (**a**) were calculated by resampling 1,000 times. The scale bars indicate substitutions per site. *1: The gene with accession No. FQ660021.1 in panel (a) was obtained from a polycycle aromatic hydrocarbon (PAH)-contaminated soil sample in a mitigated wetland.

### Universal association of COTS27 with COTS throughout the Indo-Pacific Ocean

The presence of COTS27 or COTS27-like bacteria in COTS individuals inhabiting various geographic regions was determined in a PCR assay designed to amplify a specific 261 bp fragment of the COTS27 16S rRNA gene. PCR products were obtained from all 195 COTS individuals that were collected at 15 locations throughout the Indo-Pacific Ocean comprising three known species of COTS (**Figs. 1a** and **c**). The sequencing of the PCR products from 53 randomly selected individuals confirmed the presence of COTS27 or very close relatives. The ML tree based on these 261 bp sequences (**Suppl. fig. S4**) revealed that all sequences formed a tight cluster with the six COTS27 sequences from the abovementioned phylogenetic analysis and with those obtained from the genome reconstruction described below. However, the sequences from the Israeli COTS population (Red Sea species) formed a clade separate from those of the Indo-Pacific populations from the northern Indian Ocean or Pacific Oceans. Among the northern Indo-Pacific species, only one single-nucleotide polymorphism (SNP) was detected in one sequence obtained from Japan (Wakayama C29 adult JPN; **Suppl. fig. S4**). These results indicate the universal association of COTS27 with the Indo-Pacific COTS species.

### Localization and biofilm-like structure formation of COTS27 in subcuticular spaces across the body surface of COTS

We observed the localization of COTS27 in COTS tissues using fluorescence *in situ* hybridization (FISH), as demonstrated by the binding of the general bacterial probe and a COTS27-specific probe that we designed (the binding signals on the COTS central disc spines are shown in **Fig. 4**). COTS27 was consistently present in the subcuticular spaces on both the aboral side (**Figs. 5a-d**; spines of the discs and arms, dermal papulae, and pedicellariae, see **Fig. 1b** and **Suppl. fig. S5** for their anatomical locations and structures) and oral side (**Figs. 5e**; the stems of the tube feet) and the pattern was similar for all three COTS individuals (**Figs. 4** and **5**). No COTS27 signal or any other bacterial signals were detected in the pyloric caeca and gonads (**Fig. 5g-h**). Likewise, no COTS27 was found in the pyloric stomachs, although numerous cyanobacteria-like bacteria were detected (**Fig. 5f**).

**Fig. 4.**
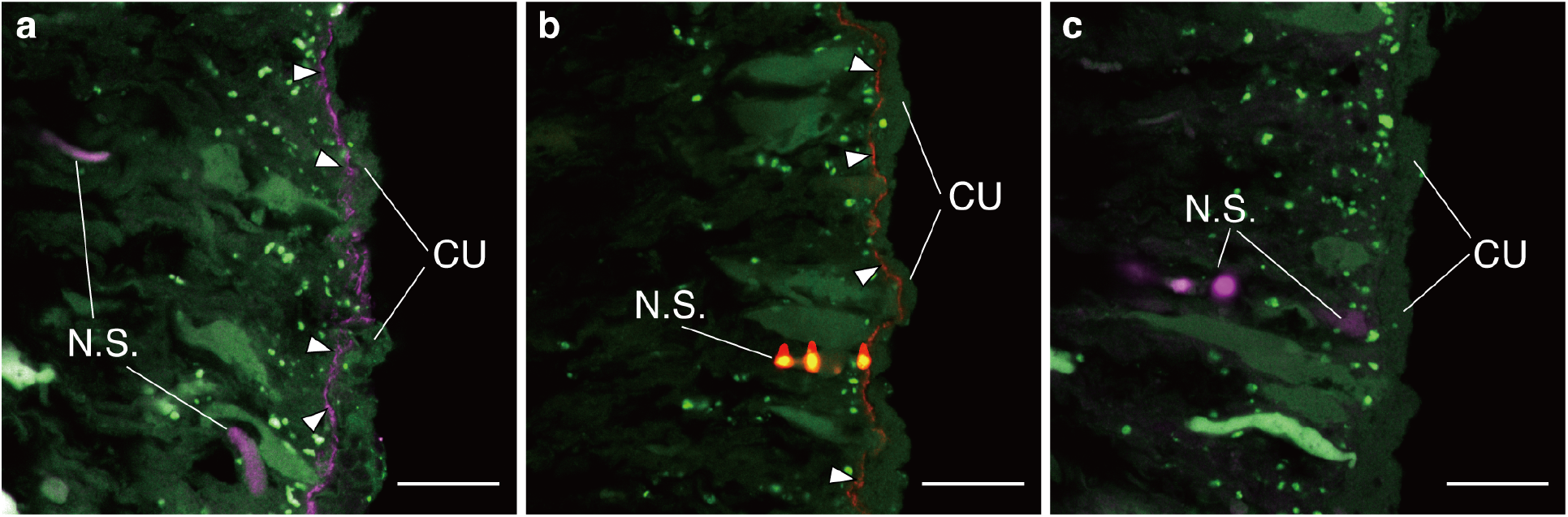
FISH analysis of three serial sections of a COTS disc spine. Each section was hybridized with the EUB338mix (**a,** purple; a general probe for bacteria), COTSsymb (**b**, red; a COTS27-specific probe), or Non338 (**c**, purple; a negative control to detect non-specific binding) probes. The probes were labelled with Cy3 in all panels and coloured with purple in panels **a** and **c** and with red in panel **b**. The green signals are tissue-derived autofluorescence. The arrowheads in panels **a** and **b** indicate layer-like signals from the general probe for bacteria (**a**) and the COTS27-specific probe (**b**). N.S. and CU indicate regions with non-specific binding and the outer cuticle complex, respectively. The scale bars represent 20 μm (**a-c**).

**Fig. 5.**
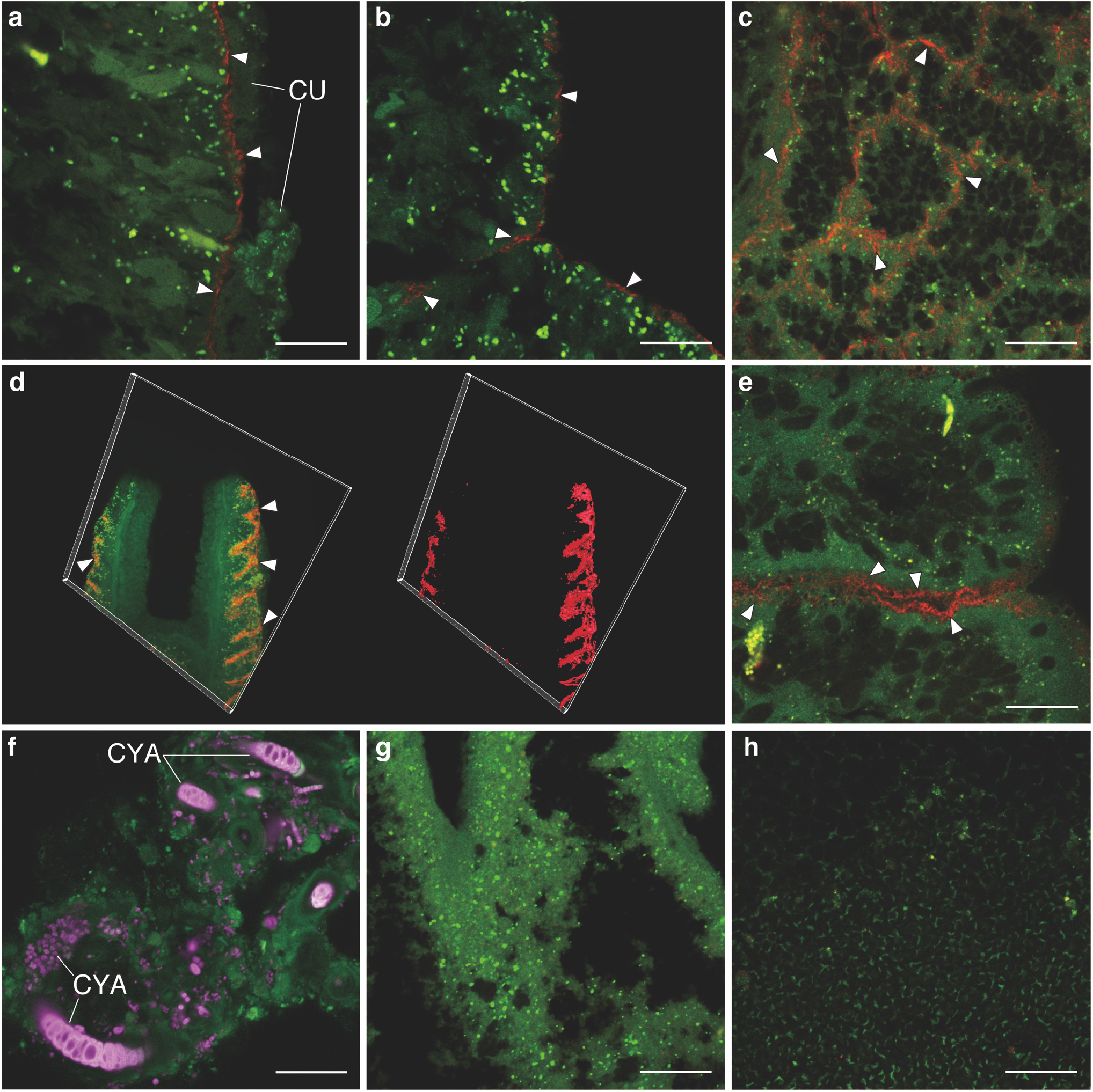
Visualization of the COTS27 cells in different body parts of COTS using FISH. COTS27 cells (red) residing in the subcuticular spaces of the body walls were detected with COTSsymb, a COTS27-specific probe, in the tips (**a**) and bases (**b**) of aboral spines on the discs and arms, respectively, dermal papula (**c**), pedicellariae on the aboral side (**d**; 3D image [left] and 3D rendering image [right]), and tube feet (**e**). Many non-COTS27 bacteria (purple) were detected in the pyloric stomachs (**f**) using the EUB338mix probe. No visible bacteria were detected in the pyloric caeca (**g**) and gonads (**h**) in this study (the images were obtained applying the COTS27-specific probe). The arrowheads indicate signals from COTS27. The green signals are tissue-derived autofluorescence. CU: outer cuticle complex; CYA: cyanobacteria-like cells. Scale bars (**a–c** and **e–h**) indicate 20 μm. The 3D image in panel (**d**) was taken with an original objective of x40.

In the cross-sections, COTS27 displayed continuous layer-like signals (**Figs. 4b, 5a-c** and **5e**), although a patchy distribution was also occasionally observed. Furthermore, three-dimensional (3D) images showed that COTS27 formed a biofilm-like structure on the epidermis of the pedicellariae (**Fig. 5d**). These observations indicate that COTS27 is an SCB that covers nearly all the surface area (the epidermis) of COTS by forming a biofilm-like structure. COTS27 cells appear to have filamentous or long rod-like shapes (**Figs. 5c** and **e**), but different approaches such as electron microscopy are required to accurately determine their cell morphology.

### Reconstruction of the COTS27 chromosome

We have not yet succeeded in isolation of COTS27, but were able to reconstruct the chromosome sequence of COTS27 from the hologenome sequences, which contained sequences derived from the host genome and the associated microbes (**Suppl. table S5**), of a COTS sample collected in Miyazaki (**Fig. 6** and **Suppl. table S6**), with 90.66% completeness and 0.26% contamination, as evaluated by CheckM [24]. The structural accuracy of the chromosome was validated based on the physical coverage of the 15 kbp-mate-pairs (**Suppl. Materials and Methods Fig. 1**), and the circular structure was also confirmed using PCR and Sanger sequencing. We also tested other assembly pipelines consisting of removal of reads from the host genome, metagenome assemblers, and binning tools; however, none generated a higher-quality genome of COTS27 nor an obvious chromosome of a different bacterium (**Suppl. Materials and Methods table 1**). Although 23 gaps remained in the final assembly, all were derived from tandem repeats in the genic regions, and the estimated gap sizes were less than 28 bp. The COTS27 chromosome was 2,684,921 bp in length, with a 39.6% average GC-content, and contained 1,650 protein-coding genes, three rRNA genes, and 35 tRNA genes. No transposable elements or prophages were detected. The 1,650 protein-coding genes included one giant gene (53,043 bp in length; COTS27_01023), but its function is currently unpredicted (see **Suppl. materials and methods** for the details). Among the three rRNA genes, the 16S rRNA gene was located separately from the 23S and 5S rRNA genes. The 35 tRNA genes covered all 20 basic amino acids. Phylogenetic analysis using the sequences of 43 conserved marker genes with 5,656 reference bacterial and archaeal genomes placed COTS27 in the phylum *Spirochaetes* (**Fig. 3b**), supporting the results of the 16S rRNA sequence-based analysis (**Fig. 3a**).

**Fig. 6.**
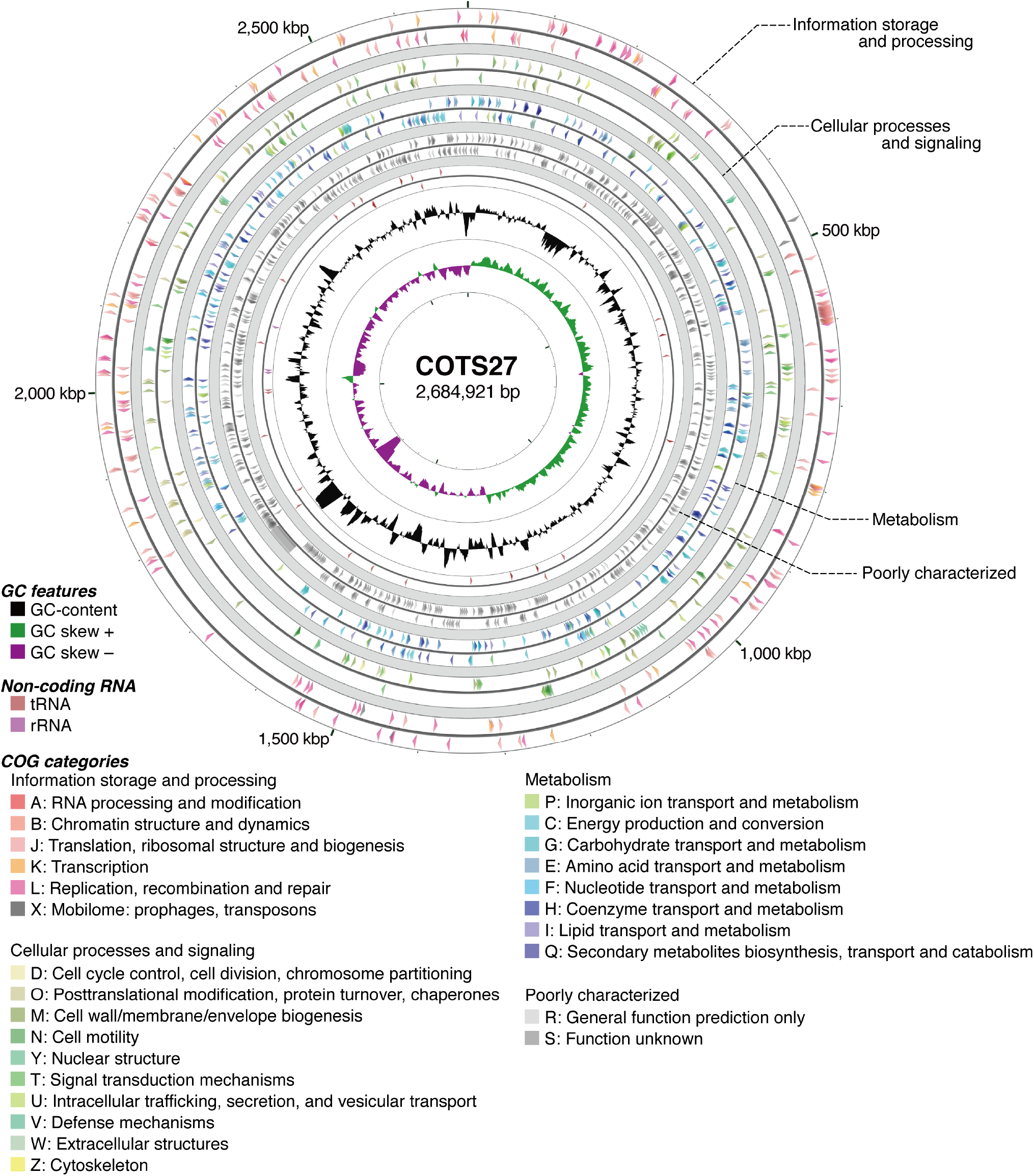
Circular view of the COTS27 chromosome. From the outside to the centre, each circle indicates forward strand CDSs; reverse strand CDSs; forward strand tRNA and rRNA genes; reverse strand tRNA and rRNA genes; GC-content; and GC skew. The CDSs were coloured according to the COG functional category of each CDS. The circular maps were created using CGView Server and the designations were then superposed manually.

### Biological features of COTS27 inferred from the gene repertoire

In the Clusters of Orthologous Groups (COG) functional category-based principal component analysis of COTS27 performed using 716 high-quality *Spirochaetes* genomes obtained from the IMG database [25], COTS27 was placed in a distinct position with regard to all *Spirochaetes* (**Suppl. fig. S6**). This indicates potential biological features unique to COTS27. Subsequently, we performed a Kyoto Encyclopedia of Genes and Genomes (KEGG) pathway analysis to obtain basic information on the biology of COTS27 (**Suppl. table S7**). Complete or near-complete biosynthesis pathways for 18 of the 20 basic amino acids were identified, excluding those for asparagine and aspartic acid. Although the guanine ribonucleotide biosynthesis pathway was not complete (one block missing), all other nucleotide biosynthesis pathways were detected. For vitamin and cofactor biosynthesis, the complete biosynthetic pathways of nicotinamide adenine dinucleotide (NAD), coenzyme A, and riboflavin and the C1-unit interconversion were identified. Pathways for fatty acid biosynthesis, beta-oxidation, and phosphatidylethanolamine biosynthesis were also detected. The conservation of these metabolic pathways suggests that COTS27 is not strongly metabolically dependent on the host COTS.

Regarding energy production, COTS27 exhibited the complete glycolysis pathway and TCA cycle. Genes for succinate dehydrogenase, cytochrome c oxidase, and F-type ATPase were also identified; however, no genes for NADH dehydrogenase were detected. Instead, COTS27 presented an operon encoding a sodium-pumping NADH:ubiquinone oxidoreductase (Na^+^-NQR) (**Suppl. fig. S7**).

Consistent with the general characteristics of *Spirochaetes*, which are generally Gram-negative, helical or spiral-shaped, and motile, with periplasmic flagella [26], COTS27 contained sets of genes for the biosynthesis of DAP-type peptidoglycan, lipopolysaccharide and phosphatidylethanolamine. While a set of genes for flagellar biosynthesis was identified, no gene for chemotaxis, such as genes encoding methyl-accepting chemotaxis proteins and chemotaxis-related signal transduction conponents, was detected.

## Discussion

We identified a single bacterium that forms a subcuticular biofilm-like structure in Indo-Pacific COTS although we can not completely exclude a possibility that the signals detected in FISH analyses included some false-positive signals (because of only *in silico* verification). This bacterium is universally present and numerically dominant and likely represents a previously undefined order within the phylum *Spirochaetes*. The universal association of COTS27 with the Indo-Pacific COTS species suggests a long history of the COTS-COTS27 association. COTS are thought to have allopatrically diverged into four species during the Pliocene-Early Pleistocene (1.95–3.65 Myr ago) in the Indo-Pacific Ocean [27]. Therefore, the association of COTS27 with at least three of the four extant COTS species (data for the fourth species are currently not available) implies that the mutual relationship between COTS and COTS27 emerged prior to the Pliocene or Early Pleistocene eras. This hypothesis is supported by the finding that COTS27 from the northern Red Sea (forming a different cluster from the Indo-Pacific regions; **Suppl. fig. S4**) was notably different from COTS27 from other regions. Additional comparative genomic analyses of the Indo-Pacific COTS and COTS27 from different geographic regions would provide more detailed insights into the possible co-evolutionary history. In addition, further studies linking environmental conditions with COTS27 abundance and microbial composition, will help to understand the ecological roles of COTS27.

Regarding the evolutionary and functional aspects of COTS27 and its association with COTS, we obtained two key findings from the genome sequence analysis: 1) the presence of Na^+^-NQR and 2) the selective loss of chemotaxis genes. 1) Some bacteria living in Na^+^-rich environments (e.g., marine or intercellular environments) exhibit an Na^+^-NQR that oxidizes NADH to NAD^+^ and pumps Na^+^ out of cells, thus functioning in the respiratory chain and in the maintenance of intercellular homeostasis in Na^+^-rich environments [28]. In line with these features of Na^+^-NQR, only one genome out of the 716 high-quality *Spirochaetes* genomes (above), which was also reconstituted from the metagenome sequences of a seawater sample (*Spirochaetaceae* bacterium NP120, IMG Genome ID: 2509276057), contained the Na^+^-NQR operon. The acquisition of Na^+^-NQR may represent one of the mechanisms responsible for the adaptation of COT27 to marine environments. 2) Besides, the selective loss of chemotaxis genes is very unusual in *Spirochaetes;* most of the high-quality *Spirochaetes* genomes mentioned above (>98%) contained gene sets for both flagellar biosynthesis and chemotaxis. The remaining genomes, such as those from the genus *Sphaerochaeta*, lack genes for both flagellar biosynthesis and chemotaxis, suggesting that chemotaxis genes have been selectively lost from the COTS27 genome. It has been proposed that the active migration and colonization by symbionts through motility and chemotaxis are often required for the acquisition of microbial partners by host organisms from environments [29]. However, the selected lost chemotaxis genes appear to represent a specific adaptation strategy of COT27 as an SCB. COTS27 may require flagella to spread and stably and widely colonize subcuticular spaces, but chemotaxis is no longer required after specially adapting to the subcuticular spaces of COTS. These findings are informing because these features are likely linked to the adaptation to marine environments and evolution as an extracellular endosymbiont in the subcuticular space, respectively. Additional genome sequences of marine spirochetes are required to verify this hypothesis and elucidate the evolutionary and functional aspects of the COTS27-COTS association.

Our study implied that COTS27 as SCB forms a nearly mono-specific symbiotic relationship with COTS, at least with the Indo-Pacific COTS. SCB, however, are commonly found in echinoderm fauna [13,19] and have been classified into three morphotypes [13,15,16]. Among these morphotypes, COTS27 most likely belongs to the SCB Type 2, which exhibits a long spiral shape and is commonly found among all five echinoderm classes [15,30]. Jackson et al. suggested the presence of a highly dominant *Spirochaetae* in the hard tissues (including the body wall) of some starfish species in the United States and Australia [31]. Such a wide distribution of spirochetes or spirochete-like bacteria in echinoderms suggests that many echinoderms may have established symbiotic relationships with marine spirochetes that are similar to that between COTS27 and COTS. Further explorations of SCB in a wider range of echinoderms would provide more detailed insights into the association between echinoderm hosts and marine spirochetes.

The outer body surfaces of marine organisms often represent a highly active interface between an organism (host) and the surrounding marine environment regarding aspects such as light exposure, gas exchange, nutrient uptake and interactions with other fouling organisms, consumers, and pathogens [32]. The presence of SCB among different echinoderms has been reviewed in different bacterial taxa [13,18,19]. Although it is also plausible that SCB play hypothetical role in their interactions with the host such as nutrition transfer [9,33], or antibiotics production [14,34], or even that their presence may be vestigial, remaining as leftovers from a previously mutualistic partnership [15], the physiological and potential ecological roles of SCB are largely unclear and remain unexplored. The universal and nearly mono-specific association of COTS27 with COTS would provide an ideal model system for further exploring the roles of SCB as well as symbiont-host-interactions in marine invertebrates. Moreover, COTS27 could be used as an environmental marker to monitor and/or predict population outbreaks of COTS.

## Conclusions

Despite the fact that the 205 COTS individuals utilized in our current analyses were collected over a 13-year period (2004–2017) and from 17 different locations across the Indo-Pacific, the COTS27 association remained exceptionally ubiquitous both spatially and temporally. Additionally, it is likely that COTS hosted COTS27 as an extracellular endosymbiont for more than 2 million years before allopatric speciation occurred during the Pliocene-Early Pleistocene suggesting a strong association. COTS27 is likely an extracellular endosymbiotic bacteria strongly associated with COTS as an SCB. COTS27 also would acquire the Na^+^-NQR system for adapting to marine environment since the speciation within phylum *Spirochaetes*. The lack of chemotaxis genes in COTS27 would physiologically associate with optimization in subcuticular space of COTS. Although the functional role of COTS27 as an SCB is still unclear, this close relationship and chromosome genome information of COTS27 described here will significantly contribute to testing the hypotheses of symbiotic function in SCB, and may also provide as a model system for studying endosymbionts in marine invertebrates more broadly.

## Materials and Methods

### Sample collection and preparation for DNA analyses and histology

We collected 205 individual COTS from 17 locations throughout the Indo-Pacific collected over a 13 year period (2004-2017; **Fig. 1a**, and **Suppl. table S8**). For 16S rRNA metabarcoding, six individuals were collected in Okinawa and Miyazaki (three from each location) in Japan (**Fig.1a**). The specimens were dissected to facilitate sampling from different body parts (7-8 body parts; **Fig. 1b,c**) which was done in triplicate or duplicate (**Suppl. table S1**). Seawater samples were also collected in triplicate from each of the last two locations. Consequently, 130 DNA samples were prepared from the six COTS individuals (**Suppl. table S1**) and six seawater DNA samples and used for the metabarcoding analysis. The tube foot DNA samples from five of the six individuals were used to determine the full-length 16S rRNA gene sequence of the dominant OTU 1. DNA samples prepared from the tube feet of 195 individuals collected from 15 geographic locations were used to examine the presence of COTS27 (the dominant OTU 1) in three species of COTS [35] (**Fig. 1a, c,** and **Suppl. table S8**). DNA from the tube feet of one individual collected in Miyazaki (Japan) was used for the COTS hologenome sequencing (**Suppl. table S8**). [36]. Note that the experiment was designed to capture both the host and an associated microbiome genomes as a hologenome. Samples of six different body parts (**Fig. 1b, c**) were prepared from the remaining other three individuals collected in Miyazaki for the FISH analyses.

We provide a more detailed description of the above collection and preparation methods in the **Suppl. materials and methods**.

### 16S rRNA metabarcoding

16S rRNA amplicon libraries (V4 region) were prepared as previously described [37,38] using the primers listed in **Suppl. table S9** and **Suppl. fig. S8** and subjected to paired-end (PE) sequencing (2 x 300 bp) using the Illumina MiSeq platform. In total, 130 DNA samples including the samples collected from 7-8 body parts of six COTS individuals (**Suppl. table S1**), and six seawater DNA samples were analysed. The obtained PE sequences were processed using software USEARCH v8.1.1861 [39] and MOTHUR v.1.36.1 [40] software for the merging, the filtering, the OTU clustering, and the taxonomic assignment of the sequence (see more detail in **Suppl. materials and methods)**. Finally, a total of 1,535,904 sequences were assigned to bacteria. The others were assigned to eukaryotes (55,377 reads containing 74.7% of COTS genes), Archaea (12,686 reads), chloroplasts (19,715 reads), or unknown origins (476,795 reads containing 99.4% of COTS genes), and were excluded from our study (**Suppl. table S2**).

### Phylogenetic analysis of OTU 1 using the full-length 16S rRNA gene sequence

Full-length 16S rRNA gene sequences of OTU 1 were obtained from each tube foot of five COTS individuals using a specific primer set for OTU 1 that was designed in this study (**Suppl. materials and methods**, **Suppl. table S9** and **Suppl. fig. S8**). The sequences were used to reconstruct the phylogenetic tree using the maximum likelihood (ML) method (**Suppl. materials and methods**). As OTU 1 was revealed to represent a unique clade of bacteria present in COTS, we hereafter refer to this bacterium as COTS27.

### PCR screening and sequencing of COTS27

In total 195 COTS individuals were screened for the presence of COTS27 on their tube feet by PCR using primers that were designed to specifically amplify a 261 bp fragment of the 16S rRNA gene (**Suppl. table S9** and **Suppl. fig. S8**). The PCR products obtained from 53 randomly selected samples from all COTS27-positive samples (n=195) were sequenced and used for phylogenetic reconstruction (**Suppl. materials and methods**).

### Fluorescence in situ hybridization (FISH)

The FISH experiments were performed on three serial sections (thickness of 5 μm) from the six body parts of the three individuals (**Fig. 1b,c**) as previously described [41]. FISH was performed separately with three different probes: COTS27-specific oligonucleotide probe (COTSsymb; for more detail of the probe design, see in **Suppl. materials and methods, Suppl. table S9** and **Suppl. fig. S8**), a Eubacterial probe (EUB338mix [42]), and a nonsense probe (Non338 [43]). Bacterial localization was observed using a confocal laser scanning microscope (LSM 550; Zeiss, Germany) (see more detail in **Suppl. materials and methods).** In addition, we reconstructed three-dimensional (3D) structures from thick sections (thickness: 50 μm) of the disc spines using a confocal laser scanning microscope (LSM770; Zeiss, Germany),

### Reconstitution of the COTS27 chromosome from the hologenome sequences of a COTS sample

Two PE libraries and six mate-pair libraries from the tube foot of one individual were prepared and sequenced using Illumina HiSeq 2500 sequencers. *De novo* assembly was performed using Platanus v. 1.2.3 [44]. To identify the COTS27-derived sequences in the hologenome assembly, scaffolds with a high coverage depth (≥×200), which probably reflected the high abundance of the species, were selected. The average coverage depth for all scaffolds was ×130, and it was assumed that most of the scaffolds from the host COTS genome exhibted coverage depths <× 200. The longest scaffold, which was identified as the COTS27 chromosome, was closed by Sanger sequencing, and an alternative assembly was obtained using Platanus-allee v. 2.0.0 [45]. A circular view of the COTS27 chromosome was generated using the CGView Server [46] with manual processing. The completeness of the final assembly was evaluated using CheckM v. 1.0.11 [24], and the structural accuracy of the assembly was validated based on the physical coverage of the 15 kbp-mate-pairs (See Suppl. materials and methods for the details). We also tested other assembly pipelines consisting of removal of reads from the host genome, metagenome assemblers, and binning tools (See Suppl. materials and methods for details).

### Gene prediction and functional annotation

Protein-coding sequences (CDSs) were predicted by using PROKKA v. 1.12 [47], followed by manual curation. For functional annotation, Clusters of Orthologous Groups (COG) were assigned by querying the CDSs against the Conserved Domain Database (CDD) with COG position-specific scoring matrices (PSSMs) using RPS-BLAST. Additionally, K numbers of Kyoto Encyclopedia of Genes and Genomes (KEGG) were assigned to each CDS; BlastKOALA [48] and KofamKOALA [49] were used to perform searches in the KEGG GENES and KOfam databases, respectively.

### Principal component analysis and phylogenetic analysis based on the genome sequences

Principal component analysis was performed based on the compositions of the COG functional categories. The genome sequences of the *Spirochaetes* bacteria were retrieved from the DOE-JGI IMG database, and 716 high-quality genomes (completeness >90% and contamination <5% as evaluated by CheckM v. 1.0.11 [24]) were retained (see **Suppl. materials and methods** for the details). Whole-genome sequence-based phylogenetic analysis was performed using CheckM to obtain the ML tree of COTS27 and 5,656 bacterial and archaeal genomes based on the sequences of 43 conserved marker genes. The tree was visualized using FigTree v. 1.4.3 (http://tree.bio.ed.ac.uk/software/figtree/).

## List of abbreviations

CDD: Conserved Domain Database
CDSs: Protein-coding sequences
COG: Clusters of Orthologous Groups
COTS: Crown-of-thorns starfish
FISH: Fluorescence *in situ* hybridization
KEGG: Kyoto Encyclopedia of Genes and Genomes
ML: Maximum likelihood
NAD: Nicotinamide adenine dinucleotide
Na+-NQ: Sodium-pumping NADH:ubiquinone oxidoreductase
OTU: Operational taxonomic unit
SCB: Subcuticular bacteria

## Declarations

### • Ethics approval and consent to participate

Not Applicable

### • Consent for publication

Not applicable

### • Availability of data and material

All sequences produced for this study have been deposited in the DDBJ under BioProject accession number PRJDB4009 for the 16S metabarcoding and COTS27 chromosome data and accession numbers LC490103 - LC490107 and LC495323 – LC495375 for the 16S rRNA gene sequence-based phylogeny.

### • Authors’ contribution

N.W., H.Y., T.I., T. H, and N.Y. conceived the research idea and designed the experiments. N.W., H.Y., Y. H, H. S, Z.F., O. B, G.E., N.T., and N.Y. conducted the field sampling. N.W., Y.G., and N.Y. performed the 16S rRNA metabarcoding. N.W. and N.Y. performed the phylogenetic analysis using the full-length 16S rRNA gene. N.W. H. S, Z.F., OB, N.T., and N.Y. performed the PCR screening and sequencing of the PCR products. N.W. conducted the FISH experiments. H.Y., R.K., D.Y., and A.T. conducted the sequencing and analysis of the COTS27 genome. N.W., H.Y., T.I., T.H, and N.Y. made major contributions to the manuscript writing and figure making. R.K., Y.G., Y.O., S.T., H.S., Z.F., O.B., and G.E. contributed to writing and editing the manuscript. All authors critically reviewed, revised and ultimately approved this final version.

## • Acknowledgements

We thank members of Yasuda Laboratory and Atsushi Toyoda Laboratory, J. Suzuki, H. Kagawa, S. Nagai, Y. R. Kubota, Y. Uchida, K. Sakai, T. Miki, C. Shinzato, L. Høj, and D. Bourne for their logistical and technical support. We also thank M. Timmers, O. Ben-Zvi, N. Phongsuwan, S. Ayalon, the Seaver Institute, and the Interuniversity Institute for Marine Sciences of Eilat for local sampling and analysis.

