## Supplementary material for "A universal subcuticular bacterial symbiont of a coral predator, the crown-of-thorns starfish, in the Indo-Pacific": Suppl. M&M

### Sample collections and preparations

#### 1) 16S rRNA metabarcoding analysis

Six adult crown-of-thorns starfishes (COTSs) were used for 16S rRNA metabarcoding, three collected in Okinawa (approx. 2 m depth) in Jul. 2017 and three in Miyazaki (approx.7 m depth) in Nov. 2017 (**Suppl. table S8**). The individuals were dissected and samples collected from different body parts (7-8 body parts; disc spines [top and base], arm spines [top and base], ambulacral spines [top and base for Okinawa, or the whole spine for Miyazaki], tube feet, and pyloric stomachs, **Fig. 1**). Each body part sample was mainly prepared in triplicate but others in duplicate (**Suppl. table S1**). One liter of sea water was also collected at each location at same time and depth (three samples at each location), and filtered on a Sterivex-GP 0.22 µm filter (Millipore, USA). A total of 136 samples of body parts and seawater filters were stored at -20˚C until DNA extraction.

#### 2) Phylogenetic analysis of dominant OTU 1 using the full-length 16S rRNA gene sequence

The tube feet of five individuals (n = 2 from Okinawa and n=3 from Miyazaki) were used from the same samples of 16S rRNA metabarcoding analysis above.

#### 3) PCR screening and sequencing of COTS27

The 195 COTS individuals from other 15 sites used for PCR screening and sequencing of one dominant OTU (OTU 1; COTS27) in their tube feet included ethanol-preserved laboratory stocks, and those from our previous studies [1,2] collected between 2004 and 2017 (for more details see **Suppl. table S8**).

#### 4) Reconstitution of the COTS27 chromosome from the hologenome sequence of a COTS samples

The tube feet of one COTS individual used for hologenome sequencing was collected in Miyazaki in Aug. 2014

#### 5) DNA extractions

All DNA samples, except for the hologenome sequencing sample, were extracted using a protocol previously described [3] and dissolved in the TE (Tris-EDTA) solution for subsequent analyses. The tube feet sample for the hologenome sequencing was stored in modified CHAOS solution (4 M guanidine thiocyanate, 0.1% N-lauroyl sarcosin sodium, 10 mM Tris pH8, 0.1 M 2-mercaptoethanol) [4,5].

#### 6) Histological procedure for Fluorescence in situ hybridization (FISH)

Three adult individuals used for FISH analyses were collected in Miyazaki (Japan) in Apr. 2017 and dissected samples collected from six body parts: aboral spines from both of disc and arms, tube feet, pyloric stomach, pyloric caeca and gonads (**Fig. 1b**). Each sample was fixed immediately in 4% paraformaldehyde phosphate buffer solution (100 mM PB; Wako,

Japan) for eight hours, and stored in 50% ethanol at 4 ˚C until next procedure. The samples were rinsed with 100 mM phosphate-buffered saline (pH 7.4; NIPPON GENE, Japan) for three times, and then decalcified by the Morse's solution [6]. After the decalcification, the samples were dehydrated through ethanol gradient series (in 70%, 90% and then in abs.100%) followed by Xylene, and embedded in paraffin. Three serial sections (5-μm thickness) were prepared on ABS-coating grass slide (MATSUNAMI, Japan) for FISH procedure.

### 16S rRNA metabarcoding

The obtained PE sequences were merged using software USEARCH v8.1.1861 [7] and filtered using MOTHUR v.1.36.1 [8] to include samples meeting the following criteria: 1) read lengths between 200 and 305 bp; 2) read quality score > 27; and 3) homopolymer read length < 8 bp. From the high-quality merged read sequences, chimeric reads were eliminated using USEARCH (parameters: reference mode, RDP Gold database, and minimum division of five) in UCHIME [9] resulting in 2,100,477 merged sequences without any ambiguous bases. Finally, these sequences were clustered into operational taxonomic units (OTUs) with a cut-off value of 97% identity using UPARSE [10]. The OTUs were assigned to known taxonomic groups by mapping onto the Silva SSU v132 database (https://www.arb-silva.de/) using MOTHUR with a cut-off value of 65. The OTUs that were assigned to unknown taxa were further processed with Silva SINA [11]; the minimum identity criterion to query the sequences was set to 0.70. Rarefaction curves were generated using the R package phyloseq ver. 1.30.0 [12] in R [13].

### Phylogenetic analysis of OTU 1 using full-length 16S rRNA gene sequences

To determine the full-length 16S rRNA gene sequences of OTU 1 we first designed two specific primers, COTS_V4_R and COTS_V4R_F (**Suppl. fig. S8** and **Suppl. table S9**), based on the sequence of OTU1 using Primer-BLAST [14]. Analyses using the Silva SSU v132 database (https://www.arb-silva.de/) and TestPrime [15] confirmed that these primers can discriminate OTU 1 from other bacterial sequences.

Next, the 16S rRNA gene was amplified by PCR using two primer sets (27F / COTS_V4_R and COTS_V4R_F / 1492R(c)), and two PCR products were directly sequenced from both directions. PCR amplifications were carried out in a 10 μl reaction mixture containing 1 μl of template DNA, 3.86 μl of $dH_2O$, 0.07 μl of each primer (50 μM), and 5 μl of KAPA Taq ReadyMix (Nippon Genetics Co. Ltd, Japan). The thermocycling program consisted of an initial denaturation at 95˚C for 2 min, 40 cycles of 94˚C for 30 sec, 50˚C for 30 sec and 72˚C for 90 sec, and a final extension at 72˚C for 5 min. The two PCR products were sequenced using the Big Dye Terminator

Sequencing kit using the PCR primers on an ABI 3730 capillary sequencer (Applied Biosystems Inc., USA).

To close the sequencing gap between the two abovementioned amplicons, we designed another primer set, Microbiont F and R (**Suppl. fig. S8** and **Suppl. table S9**), and used another PCR amplification for that gap region. PCR amplifications were performed in a 10 μl reaction mixture as described above. The thermocycling was performed with an initial denaturation at 94˚C for 1 min, 40 cycles of 94˚C for 20 sec, 55˚C for 45 sec and 72˚C for 3 min, and a final extension for 10 min at 72 ˚C. The PCR products were sequenced as described above.

The sequences were assembled using ATSQ software (GENETYX, Japan) to reconstruct full-length 16S rRNA gene sequences of OTU 1. We retrieved all sequences of the phylum *Spirochaetes* available in the Living Tree Project (LTP) release 128 [16]. We also obtained additional reference sequences that clustered with OTU 1 using the Silva SSURef_NR99_128 database (https://www.arb-silva.de/). All sequences were aligned using MUSCLE [17]. Phylogenetic tree was constructed using the Maximum likelihood (ML) method with the generalized time-reversible model with gamma distribution and proportion of invariable sites, applying 1000 bootstrap replications in MEGA7 [18]. As OTU 1 formed a unique clade it was renamed COTS27.

### PCR screening and sequencing of COTS27

Two PCR primers (COTSsymb F and R; **Suppl. fig. S8** and **Suppl. table S9**) were designed using the Primer-BLAST [14] to specifically amplify a 261-bp region of the COTS27 16S rRNA genes region. The specificity of these primers to discriminate COTS27 from other bacterial sequences including other *Spirochaetes* was confirmed (Non-match in the database was verified using TestPrime [15]). An *Asteroidea*-universal primer set (hitode_16S f & r; **Suppl. table S9**) that amplifies mitochondrial 16S rRNA gene sequence was also designed and used as a positive control for PCR reactions. PCR amplifications were performed in a 10 μl reaction mixture containing 1 μl of the genomic DNA, 3.86 μl of $dH_2O$, 0.07 μl of each primer (50 μM) and 5 μl of KAPA Taq ReadyMix (Nippon Genetics Co. Ltd). In several cases, KAPA2G Robust HotStart ReadyMix (Nippon Genetics Co. Ltd, Japan) or Go Taq master mix (Takara, Japan) was used instead of KAPA Taq ReadyMix. PCR products were detected by electrophoresis on a 1% agarose gel. Selected PCR products (n=53) were sequenced using the COTSsymb F and R primers and sequencing data used for constructing phylogenetic tree as described above.

### Fluorescence in situ hybridization (FISH)

The specific probe COTSsymb (*E. coli* position 196)) for COTS27 was designed on 16S rRNA gene using the ARB software package [19] with the SSURef_NR99_128 database from ARB SILVA (https://www.arb-silva.de/) (**Suppl. fig. S8** and **Suppl. table S9**). *In silico* evaluation using the Silva SSU r138 database with the TestProbe 3.0 tool (https://www.arb-silva.de/) and in the Ribosomal Database Project (RDP) Release 11 in the PROBE MATCH tool [20] indicated a high specificity of the probe; there was no perfect-match probe binding sites (even when predicting one miss-match) with any non-target bacteria. Furthermore, the probe optimization was validated in the generated formamide dissociation curves using mathFISH tool [21].

Bacterial localization for all samples was observed using a confocal laser scanning microscope (LSM 550; Zeiss, Germany) with two channels for Cy3 fluorescence (excitation: 555 nm; emission: BP 490-635) and COTS autofluorescence (excitation: 488 nm; emission: non-filter).

In addition, the thick sections of the disc spines for the three-dimensional (3D) structures were observed using the Z-stack function (interval of 0.2 μm) of the LSM 770 confocal laser scanning microscope (Zeiss, Germany) with two channels for Cy3 fluorescence (excitation: 561 nm; emission: 561-641 nm) and COTS autofluorescence (excitation: 405 nm; emission: 410-552 nm) and reconstructed as object using the surface rendering function of Imaris software (BitplaneAG, USA).

We also verified that hybridization of the COTS symb probe did not occur in cells from an *E. coli* culture, and we applied a non-sense probe as a negative control to ensure that true COTS27 bacterial signals could be distinguished from non-specific probe binding.

### Hologenome sequencing analysis

***Reconstruction of the COTS27 chromosome sequence:*** Two paired-end (PE) libraries (insert sizes; 300 bp and 500 bp) were prepared and sequenced using the Illumina HiSeq 2500 sequencer. *De novo* assembly was performed using Platanus v. 1.2.3 [22]. To identify the COTS27-derived sequences, we extracted long scaffolds (≥5000 bp) that had higher depths of coverage (≥200×) compared to the average of all scaffolds (130×). Coverage depths were estimated using Platanus. In addition to the PE libraries, we prepared and sequenced six mate-pair (MP) libraries (insert sizes; 3, 5, 8, 10, 12, and 15 kb). Using all MP reads, additional scaffolding was performed for the potentially COTS27-derived scaffolds. As the longest scaffold was suspected as the COTS27 chromosome, several gaps in the scaffold were closed by PCR and Sanger sequencing as well as *in silico* based on the assembly results obtained using another assembler, Platanus-allee v. 2.0.0 [23]. The completeness of the obtained sequence was estimated by Check M [24].

***Trials of metagenome assembly pipelines:*** The pipelines designed for metagenome (hologenome) assembly were tried as follows:

(1) Mapping of reads to the host genome

All paired-end (PE) and mate-pair (MP) reads were mapped to the reference genomes of COTS [25] (accessions, GCF_001949145.1 and GCA_001949165.1) using Bowtie2 v. 2.3.4.3[26]. The tool was executed as the single-end mode (each file of reads was input using the option of "-U") with the option of "--very-sensitive-local". The mapping results whose identity ≥90% and alignment-coverage ≥50% were extracted.

(2) Removal of reads from the host genome

Each read pair was excluded if at least one read was mapped to the COTS genome as a sequence from the host COTS.

(3) *De novo* assembly

The resultant reads of (2) were assembled using Platanus v. 1.2.3[27], SPAdes v. 3.1.3.1[28], metaSPAdes v3.13.1[29], IDBA-UD v. 1.1.3[30] and MEGAHIT v. 1.1.3[31]. Note that the latter three assemblers were designed for metagenome data. Platanus was executed with the option of "-n 20" for "assemble" command, which means an initial coverage cutoff was 20. The other assemblers were executed with the default parameters. Only PEs were input into metaSPAdes and MEGAHIT because of the software limitations. 300-PE, 500-PE, 5k-MP, 10k-MP and 15k-MP were input into IDBA-UD because the maximum number of input libraries was five.

(4) Removal of assembled sequences from the host genome

It was assumed that sequences from the host genomic regions that were highly differentiated from the COTS reference genomes. The assembly results were searched against the NCBI nt database (release, 2019-6) using BLASTN v. 2.6.0[32] with the e-value cutoff of 1e-5. The sequences whose best-scoring hits corresponded to *Acanthaster planci* (taxonomy ID, 133434) were excluded.

(5) Binning

For each assembly result, MaxBin2 v. 2.2.6[33] was applied to construct metagenome bins (sequence sets). MaxBin2 utilizes information of coverage depth (abundance), single-copy marker genes and tetranucleotide frequencies. The coverage depths of paired-ends were calculated using BWA-MEM v. 0.7.12-r1044[34] and the program of jgi_summarize_bam_contig_depths (https://bitbucket.org/berkeleylab/metabat/downloads/metabat-static-binary-linux-x64_v2.12.1.tar.gz) with the default parameters, and the results were input into MaxBin2. The length cutoff of input assembly was set to 3000 ("-min_contig_length 3000" when MaxBin2). Sequences that were not contained in bins were discarded, and statistics were calculated for each bin.

(6) Alignment to the COTS27 chromosome

Each metagenome bin was aligned to the COTS27 chromosome we constructed using FastANI v. 1.1[35] with the default parameters. For bins of which alignments were detected, all of them indicated average nucleotide identity (ANI) $\geq$99% and alignment coverage $\geq$95%. These bins and the others were categorized into COTS27-bins and non-COTS27-bins, respectively.

(7) BLASTN search of non-COTS27 bins against the database

Non-COTS27-bins were searched against the NCBI nt database (release, 2019-6) using BLASTN v. 2.6.0 with the e-value cutoff of 1e-5. For each bin, the top-scoring hits were determined based on the sum of the bit-scores of local alignments.

***Gene prediction and functional annotation:*** Gene prediction and functional annotation were performed using PROKKA v. 1.12 [36]. Predicted genes were manually confirmed utilizing In Silico Cloning Genomic Edition v. 5 (In Silico Biology Inc., Yokohama, Japan). Annotated gene names were curated using the following sources of information: (1) Blast hit table for the National Center for Biotechnology Information (NCBI) NR database and UniProt Knowledgebase (UniProtKB) Swiss-Prot, (2) protein information in UniProtKB, (3) protein signatures detected using InterProScan v. 5.22-61.0 (Jones et al. 2014), (4) KEGG assignments of proteins by BlastKOALA v. 2.1 and KofamKOALA v. 2019-04-06, (5) operon information for *Escherichia coli* K-12 in the RegulonDB [37], and (6) membrane-related characteristics of proteins predicted using SOSUI version 1.10 [38]. COG assignments were obtained following the Joint Genome Institute (JGI) Microbial Genome Annotation Pipeline [39]. Using the COG, Position-Specific Scoring Matrices (PSSMs) obtained from Conserved Domains Database (CDD), genes were classified according to COG functional categories using RPS-BLAST (top hit, e-value cutoff of 1e-2, alignment length of at least 70% of the consensus sequence length). To obtain metabolic pathway information, K numbers were assigned using BlastKOALA and KofamKOALA. BlastKOALA was first used to search in the KEGG GENES database (selected taxonomic group, Bacteria; selected database, “species_prokaryotes”). COTS27 Genes that were not assigned by BlastKOALA were subjected to search in the KOfam database using KofamKOALA.

To predict the function of the giant gene, following 5 methods were additionally employed: (1) The protein was searched against the Uniprot (SwissProt and TrEMBL) and NCBI-nr databases using BLASTp version 2.10.0 with an e-value cutoff of 1×10-5 [40], (2) domains were examined using InterProscan version 5.22.61 [41], (3) the presence of signal peptides was predicted using SignalP version 5.0b [42], (4) the presence of transmembrane helices was predicted using TMHMM version 2.0c [43], and (5) Subcellular localization was predicted using PSORTb version 3.0.2 with selecting Gram-negative bacteria [44].

***Validation of the structural accuracy of the COTS27 chromosome:*** The structural accuracy of the COTS27 chromosome was validated based on the physical coverage (the number of spanning read pairs) of the 15 kbp-mate-pairs (nominal insert size was 15 kbp). First, the mate-pairs were re-mapped to the COTS27 chromosome using Bowtie2 v. 2.3.4.3[26]. The tool was executed as the single-end mode (each file of reads was input using the option of "-U") with the option of "--very-sensitive-local". The mapping results whose identity ≥95% and alignment-coverage ≥75% were extracted. Next, the mapped pairs with reasonable insert-sizes (ranging from 0.5×15k to 1.5×15k) were further extracted, and physical coverage was calculated for each site. Here, high physical coverage infers high structural accuracy of a sequence. The circular structure of the chromosome was validated (*i.e.,* the head and tail of the sequence were contiguous) using PCR and Sanger sequencing.

***Comparison with other Spirochetes in the IMG database:*** A total of 834 Spirochetes genomes with COG annotation assigned were obtained from the IMG database. To select high-quality genome data, the 834 genomes were evaluated by CheckM using the *Spirochaetes* gene markers, and medium- or low-quality genomes (≤90% completeness and ≥5% contamination) were eliminated according to Bowers *et al.* 2017. Finally, 716 were retained as high-quality genome data (see **Suppl. Materials and Methods Fig. 2**) and used for the comparison with COTS27. PCA was performed using COG functional categories by the prcomp command and maptools package ver. 0.9-5 [45] in R [13].

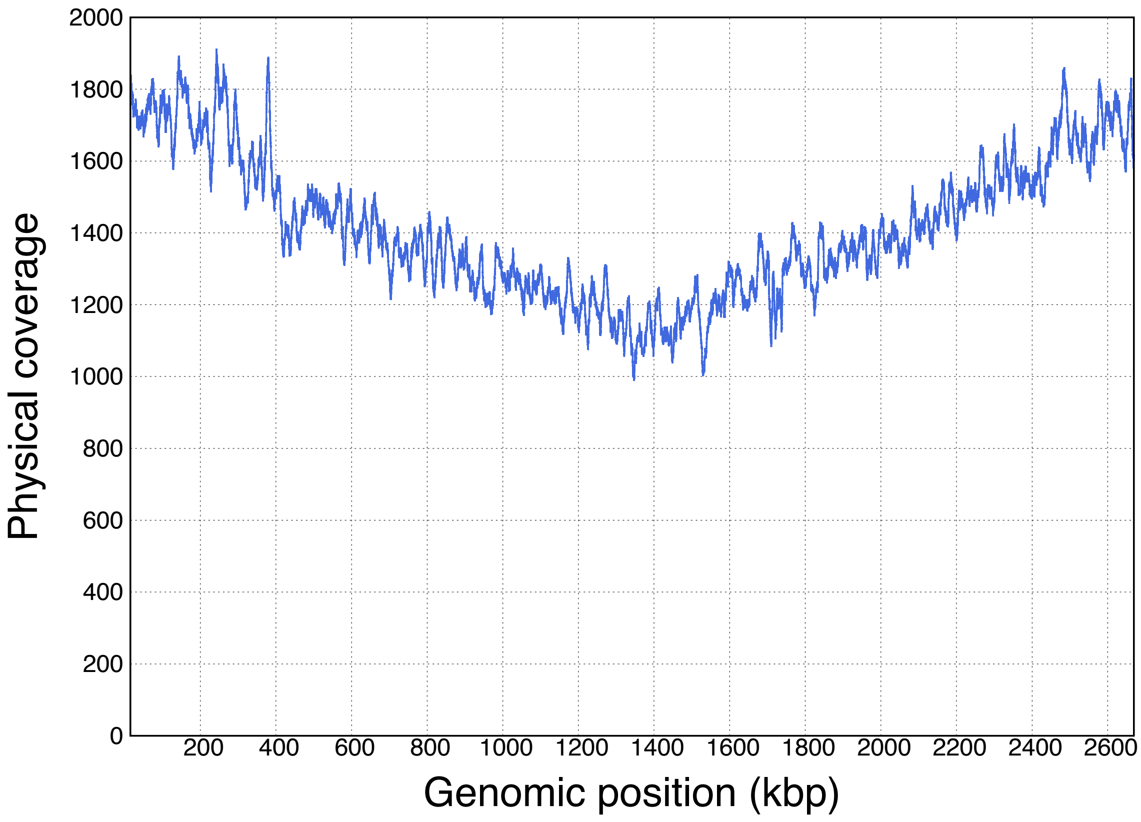


**Suppl. Materials and Methods Fig. 1 Physical coverage of 15 kbp-mate-pairs on the COTS27 chromosome**

Sites (horizontal axis) within 15 kbp from the edge are not displayed. All sites in the graph indicate physical coverages ≥988, inferring no structural error.

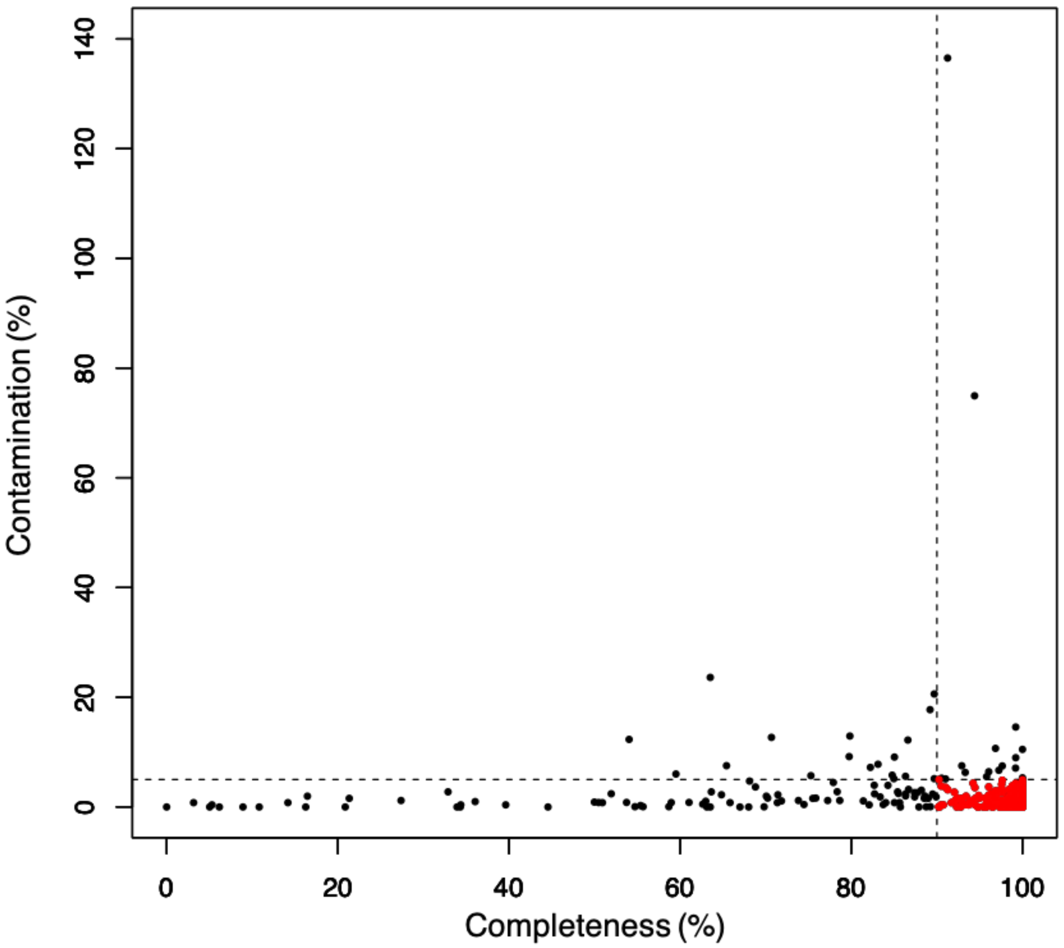


**Suppl. Materials and Methods Fig. 2 Scatter plot of CheckM assignments of 834 Spirochaetes genomes obtained from the IMG database.**

Vertical and horizontal dashed lines correspond to 90% completeness and 5 % contamination, respectively. Red dots indicate the 716 high-quality genomes showing > 90% completeness and < 5% contamination.

**Suppl. Materials and Methods table 2 Trials of metagenome assembly pipelines.**

(a) Assembly statistics. (b) COTS27-bins statistics. (c) CheckM-evaluations and alignments to the result of our pipeline. (d) Non-COTS27 bin statistics. (e) NCBI nt top hits of non-COTS27 bins.

"ANI": average nucleotide identity. Compared to our pipeline, no other pipeline indicates better assembly contiguity (maximum and N50 lengths of scaffolds and contigs) nor a much larger total size (all bins had total sizes <2.703 Mbp). For non-COTS27 bins, CheckM-completeness were both 0%. The non-COTS bins may have resulted from the low-abundance species or contamination.

(a) Assembly statistics.

| Pipeline | Total (bp) | # scaffolds | Scaffold N50 (bp) | Scaffold max (bp) | # contigs | Contig N50 (bp) | Contig max (bp) | Gap ('N') rate (%) | # bins |
|---|---|---|---|---|---|---|---|---|---|
| Our pipeline | **2,684,921** | **1** | **2,684,921** | **2,684,921** | **24** | **404,513** | **855,651** | **0.01** | **1** |
| Platanus-based | 2,702,213 | 6 | 2,669,157 | 2,669,157 | 58 | 232,230 | 652,618 | 0.44 | 1 |
| IDBA-UD-based | 5,499,917 | 210 | 69,826 | 328,741 | 238 | 68,207 | 303,592 | 0.04 | 2 |
| MEGAHIT-based | 2,677,178 | 83 | 91,525 | 220,053 | 83 | 91,525 | 220,053 | 0.00 | 2 |
| SPAdes-based | 2,935,562 | 41 | 2,498,517 | 2,498,517 | 133 | 81,898 | 327,747 | 6.95 | 2 |
| metaSPAdes-based | 2,609,088 | 56 | 151,647 | 201,380 | 120 | 46,944 | 179,544 | 0.23 | 1 |

(b) COTS27 bin statistics.

| Pipeline | Bin ID | Total (bp) | # scaffolds | Scaffold N50 (bp) | Scaffold max (bp) | # contigs | Contig N50 (bp) | Contig max (bp) | Gap ('N') rate (%) |
|---|---|---|---|---|---|---|---|---|---|
| Our pipeline | **Our-pipeline-bin1** | **2,684,921** | **1** | **2,684,921** | **2,684,921** | **24** | **404,513** | **855,651** | **0.01** |
| Platanus-based | Platanus-bin1 | 2,702,213 | 6 | 2,669,157 | 2,669,157 | 58 | 232,230 | 652,618 | 0.44 |
| IDBA-UD-based | IDBA-UD-bin1 | 2,674,748 | 57 | 165,273 | 328,741 | 85 | 106,579 | 303,592 | 0.08 |
| MEGAHIT-based | MEGAHIT-bin1 | 1,246,988 | 26 | 107,262 | 220,053 | 26 | 107,262 | 220,053 | 0.00 |
| MEGAHIT-based | MEGAHIT-bin2 | 1,430,190 | 57 | 69,950 | 201,904 | 57 | 69,950 | 201,904 | 0.00 |
| SPAdes-based | SPAdes-bin1 | 2,675,911 | 15 | 2,498,517 | 2,498,517 | 63 | 97,918 | 327,747 | 2.84 |
| metaSPAdes-based | metaSPAdes-bin1 | 2,609,088 | 56 | 151,647 | 201,380 | 120 | 46,944 | 179,544 | 0.23 |

(c) CheckM-evaluations and alignments to COTS27.

| Pipeline | Bin ID | CheckM completeness (%) | CheckM contamination (%) | ANI to COTS27(%) | Coverage of alignment to COTS27 (%) |
|---|---|---|---|---|---|
| Our pipeline | **Our-pipeline-bin1** | **88.76** | 0.00 | N/A | N/A |
| Platanus-based | Platanus-bin1 | 88.76 | 0.00 | 99.987 | 99.107 |
| IDBA-UD-based | IDBA-UD-bin1 | 88.76 | 0.00 | 99.987 | 97.688 |
| MEGAHIT-based | MEGAHIT-bin1 | 38.59 | 0.00 | 99.994 | 98.267 |
| MEGAHIT-based | MEGAHIT-bin2 | 50.23 | 0.00 | 99.996 | 95.815 |
| SPAdes-based | SPAdes-bin1 | 88.76 | 0.00 | 99.919 | 97.062 |
| metaSPAdes-based | metaSPAdes-bin1 | 88.76 | 0.00 | 99.960 | 99.289 |

(d) Non-COTS27 bin statistics

| Pipeline | Bin ID | Total (bp) | # scaffolds | Scaffold N50 (bp) | Scaffold max (bp) | # contigs | Contig N50 (bp) | Contig max (bp) | Gap ('N') rate (%) |
|---|---|---|---|---|---|---|---|---|---|
| IDBA-UD-based | IDBA-UD-bin2 | 2,825,169 | 153 | 39,665 | 146,206 | 153 | 39,665 | 146,206 | 0.00 |
| SPAdes-based | SPAdes-bin2 | 259,651 | 26 | 21,512 | 99,241 | 70 | 3,101 | 6,252 | 49.30 |

(e) NCBI nt top hits of non-COTS27 bins.

| Pipeline | Bin ID | Identity (%) | Sum alignment length (bp) | Top-hit descriptin |
|---|---|---|---|---|
| IDBA-UD-based | IDBA-UD-bin2 | 99.38 | 52,260 | Escherichia coli strain ST540, complete genome |
| SPAdes-based | SPAdes-bin2 | 99.40 | 55,072 | Escherichia coli strain ST540, complete genome |

***Reference***

45. Bivand R, Lewin-Koh N. maptools: Tools for reading and handling spatial objects. R Package Version 08. 2013;23.
