## Supplementary material for "A universal subcuticular bacterial symbiont of a coral predator, the crown-of-thorns starfish, in the Indo-Pacific": Suppl. appendix

*Appendix 1 Other relatively abundant bacteria in COTS*

Relatively abundant OTUs other than OTU 1 (COTS27) in COTS are shown in Appendix table 1. They belong to the families: *Spiroplasmataceae* (OTU 3, 5.8% average abundance in all COTS samples; OTU 4, 2.4%), *Bacillaceae* (OTU 5, 3.7%; OTU9, 1.9%), *Rhizobiaceae* (OTU 7, 2.8%), *Burkholderiaceae* (OTU 6, 2.4%), *Flavobacteriaceae* (OTU 8, 1.6%), *Francisellaceae* (OTU 11, 1.4%) and *Endozoicomonadaceae* (OTU 10, 0.8%).

**Appendix table 1** Other relatively abundant OTUs in COTS

| OTU ID, putative taxon | Top BLAST match *1; Accession (% identity) | Average abundance (%) in; Total COTS samples | Spine | Tube feet | Pyloric stomach |
|---|---|---|---|---|---|
| OTU 3, *Spiroplasmataieae* *2 | MG776020.1 (99%) | 5.79 | 2.83 | 4.02 | 23.03 |
| OTU 5, *Bacillaceae* | KF975538.1 (98%) | 3.67 | 3.33 | 1.19 | 7.97 |
| OTU 7, *Rhizobiaceae* | JQ387353.2 (99%) | 2.77 | 2.70 | 0.69 | 5.24 |
| OTU 6, *Burkholderiaceae* | MG858809.1 (99%) | 2.43 | 2.26 | 0.70 | 5.06 |
| OTU 4, *Spiroplasmataieae* *2 | MG776020.1 (90%) | 2.39 | 2.54 | 0.09 | 3.88 |
| OTU 9, *Bacillaceae* | KY989221.1 (98%) | 1.89 | 1.69 | 0.49 | 4.31 |
| OTU 8, *Flavobacteriaceae* | KU578369.1 (92%) | 1.62 | 1.37 | 2.55 | 1.98 |
| OTU 11, *Francisellaceae* | FJ202895.1 (98%) | 1.35 | 0.60 | 0.34 | 6.23 |
| OTU 10, *Endozoicomonadaceae* | AM495252.1 (98%) | 0.77 | 0.95 | 0.28 | 0.31 |

*1 Top BLAST hits in the NCBI nr/nt database (https://blast.ncbi.nlm.nih.gov/Blast.cgi?PAGE_TYPE=BlastSearch) are shown.

*2 The phylogenies of OTU 3 and OTU 4 were determined in the databases of All-species Living Tree Project and RDP, respectively using Silva SINA [1].

*Appendix 2 The members of clade I in marine spirochetes*

Among the marine spirochetes, COTS27 formed a distinct clade (clade I in **Fig. 3**) with an uncultured spirochete SRODG048 (GenBank accession No. FM995181) - obtained from Sydney rock oysters in Australia [2], an uncultured spirochete bacterium clone GHI14 (GenBank accession No. EU857763) - detected from crystalline styles of marine bivalves in North sea [3], and an uncultured marine bacterium clone Sp02sw36 (GenBank accession No. HQ241817) - obtained from the sponge *Tsitsikamma favus* in South Africa [4]. However, the 16S rRNA gene of COTS27 shares 85.9–86.9%, 86.4–86.9%, and 84.8–85.4% sequence identity with the three abovementioned clones, respectively. As these identity values are at the same level as the proposed threshold for assigning a novel bacterial family (86.5%) [5], COTS27 represents a distinct family in clade I.
