## Supplementary material for "A universal subcuticular bacterial symbiont of a coral predator, the crown-of-thorns starfish, in the Indo-Pacific": Suppl. figs

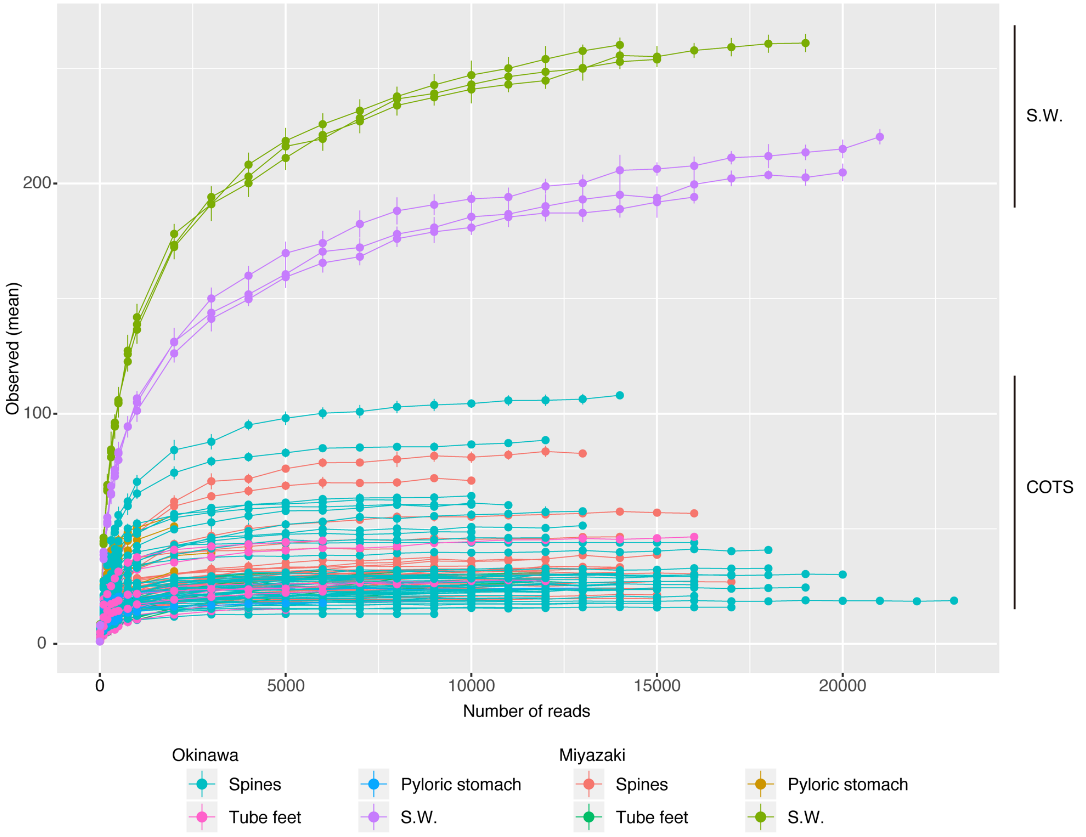


**Suppl. fig. S3 Rarefaction curves of bacterial OTUs from COTS and seawater.** Bacterial communities from the spines, tube feet, pyloric stomach, and seawater (S.W.) are shown for samples from both Okinawa and Miyazaki, Japan.

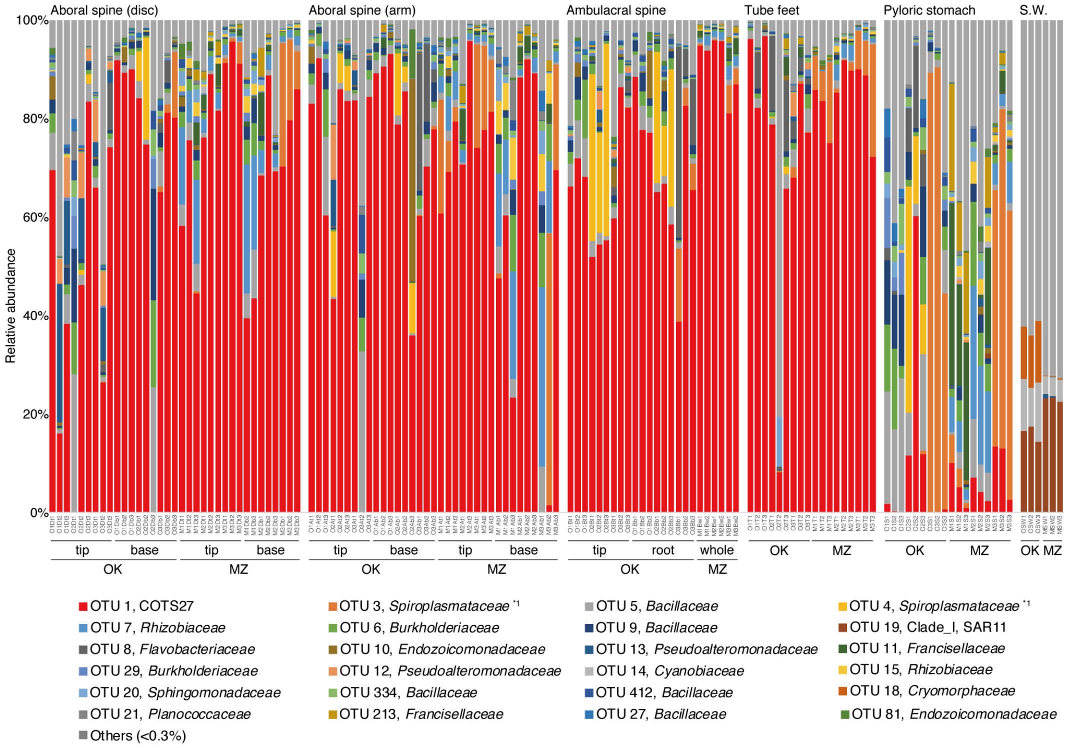


**Suppl. fig. S2 Stacked bar-plots of the relative abundance of bacterial OTUs.**

The results of family-level assignment (Silva database, cutoff ≤65) are shown. “Others” (grey) represent the sum of bacterial OTUs which occupied <0.3% of the total sequences across all samples. “S.W.” indicates seawater samples. *1: The phylogenetic statuses of OTU3 and OTU4 were determined in the databases of All-species Living Tree Project and RDP, respectively, using Silva SINA [1]. Note that OTU1 (COTS27, red) represents the most dominant bacterial OTU in most samples from the body surfaces of COTS collected at two locations in Japan (Okinawa and Miyazaki represented by OK and MZ, respectively).

1. Pruesse E, Peplies J, Glöckner FO. SINA: Accurate high-throughput multiple sequence alignment of ribosomal RNA genes. Bioinformatics. 2012;28:1823–9.

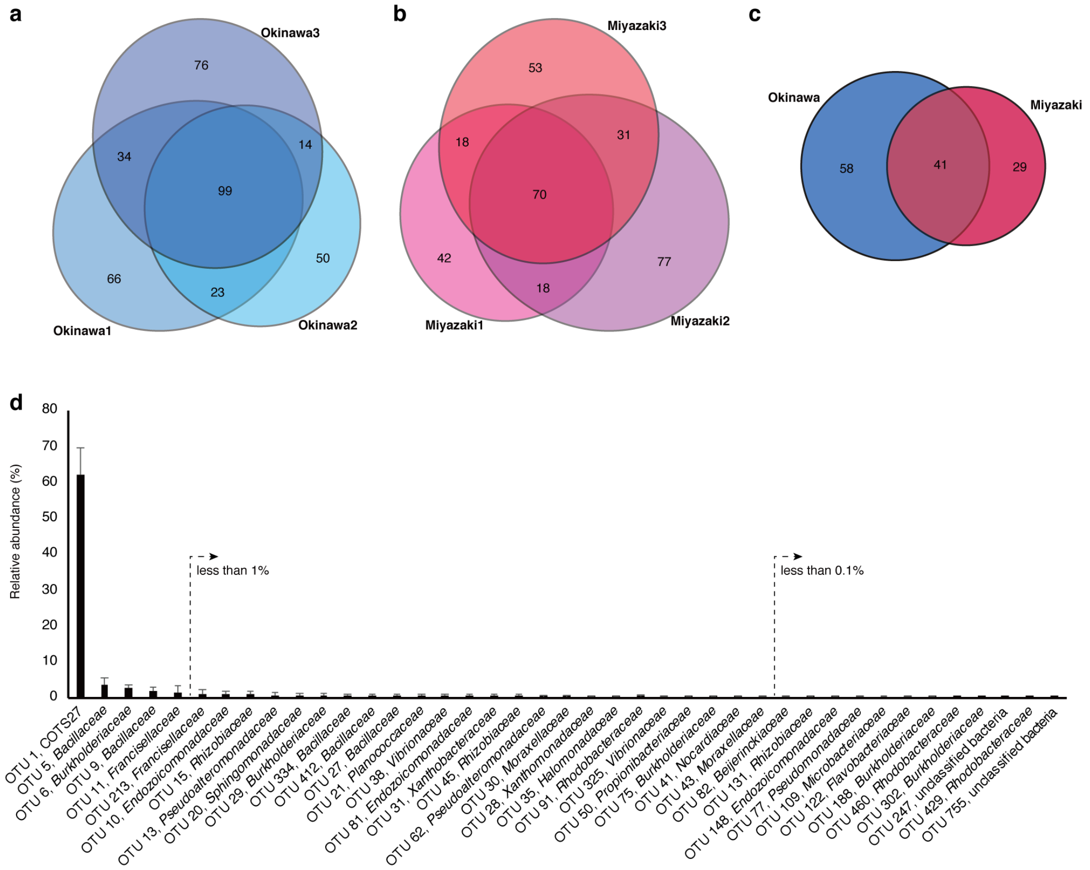


**Suppl. fig. S3 Bacterial OTUs shared by COTS individuals.**

Venn diagrams of bacterial OTUs shared by three COTS individuals collected in Okinawa (**a**) and Miyazaki (**b**), Japan. A total of 99 OTUs were shared between the Okinawan samples and 70 OTUs between the Miyazaki COTS samples. 41 OTUs were present in all COTS samples from the two locations (**c**), indicating these 41 OTUs represent the core bacteria of the COTS-associated microbiome. The relative abundance of 41 overlapping OTUs (**d**).

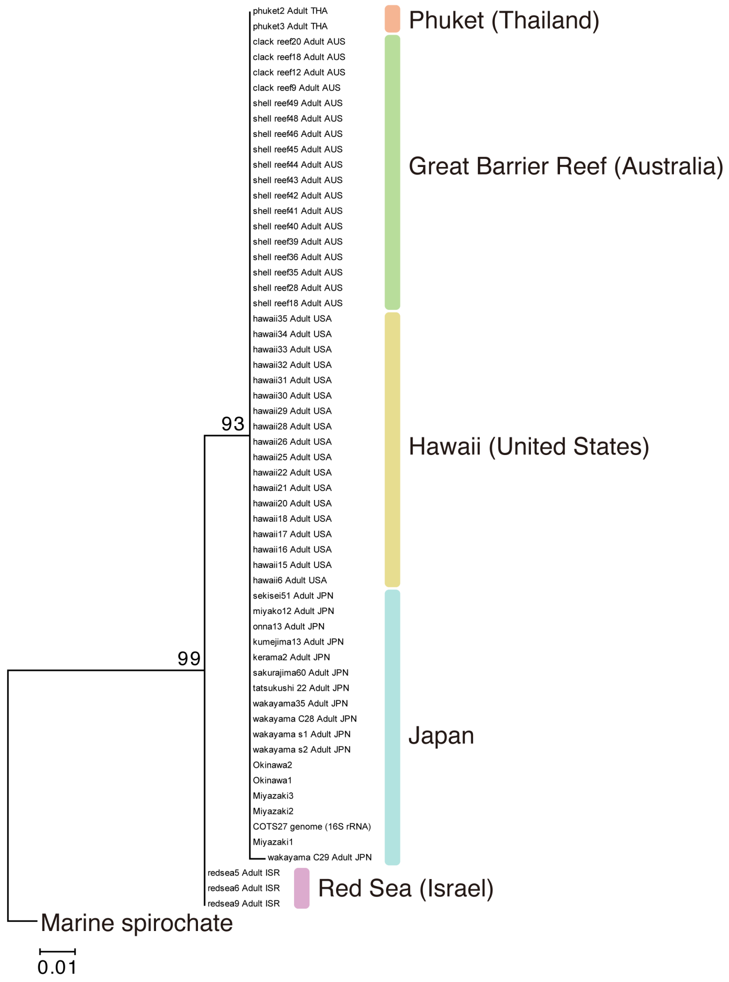


**Suppl. fig. S4 Phylogenetic analysis of COTS27 obtained from 59 COTS individuals.** Partial 16S rRNA sequences (235 bp long) of COTS27 obtained from 53 randomly selected COTS individuals (from a total of 195 PCR-positive, COTS27-specific, COTS individuals). Corresponding sequences were obtained from the five samples used for full-length 16S rRNA sequence determination and one sample used for whole genome determination, and included in the phylogenetic analysis. Thus, the reconstructing phylogenetic tree included 59 COTS27 sequences. The tree was constructed using the maximum likelihood method with 1000 bootstrap replications by MEGA7 software [2]. The tree was rooted on a member of the clade I marine spirochetes (see Fig. 3 in the main text). Note that the sequences from COTS individuals collected in Israel form a group distinct from those collected in other regions.

2. Kumar S, Stecher G, Tamura K. MEGA7: Molecular Evolutionary Genetics Analysis Version 7.0 for Bigger Datasets. Mol Biol Evol. 2016;33:1870–4.

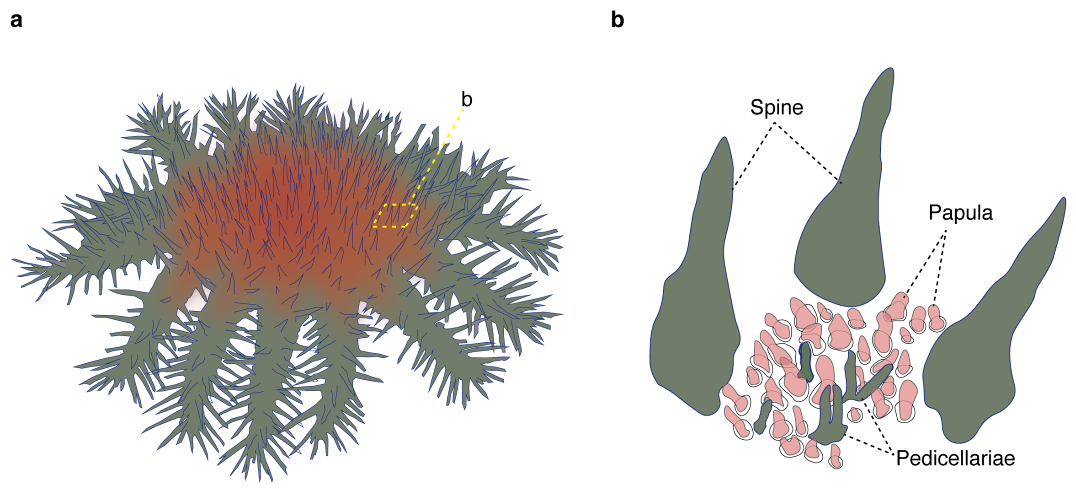


**Suppl. fig. S5 Schematic drawing showing the dermal papula (skin gill) and pedicellariae (small external appendages) between disc spines (b) on the body surface of COTS (a).**

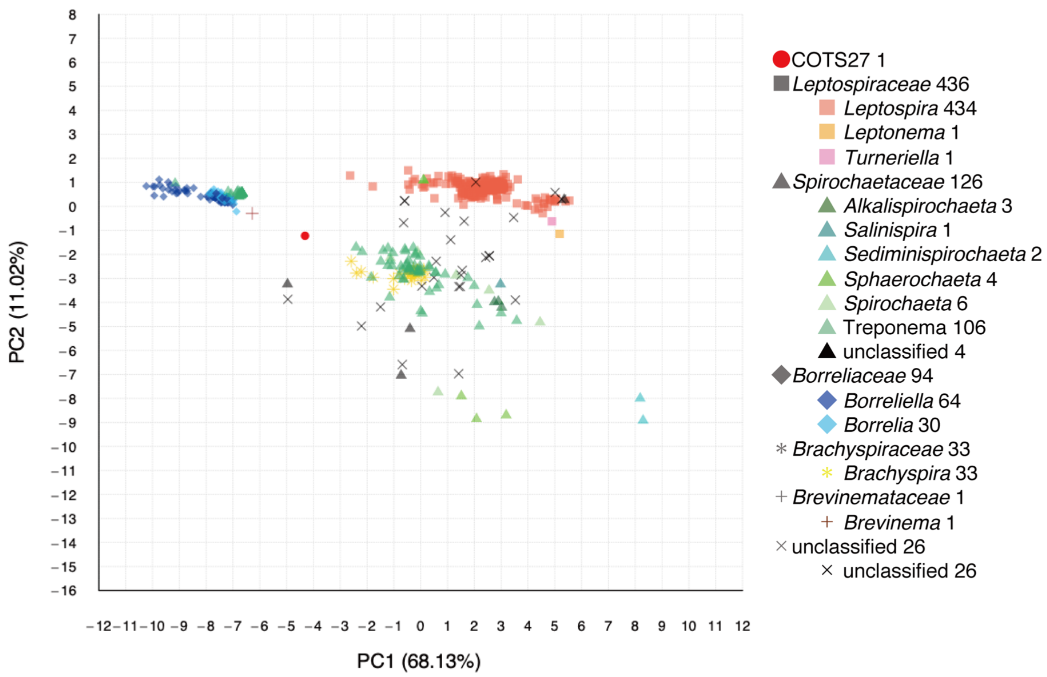


**Suppl. fig. S6 Principal component analysis (PCA) of COTS27 and 716 high-quality *Spirochaetes* genomes based on the gene abundance according to the COG functional categories.**

The red circle indicates COTS27. Different families or genera are indicated by symbols with different shapes and colors. Numbers next to family or genus names indicate the number of strains belonging to each family or genus.

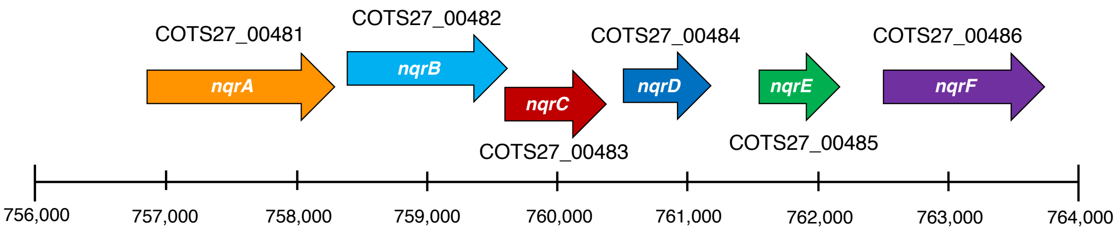


**Suppl. fig. S7 Gene organization and position of the $Na^+$-NQR genes on the COTS27 chromosome.**

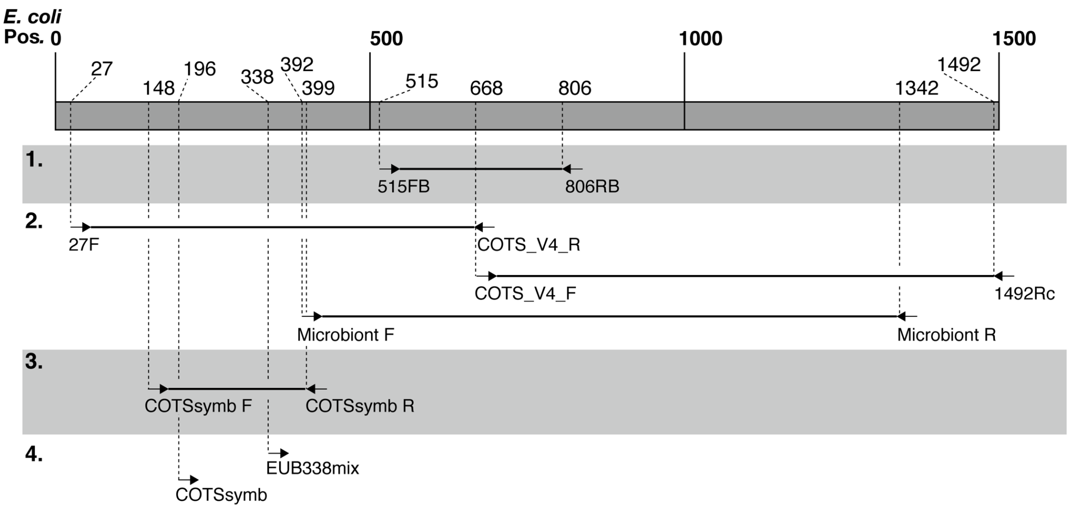


**Suppl. fig. S8 Positions of the primers and probes used in the current study for the 16S rRNA gene analysis.**

**1.** Primers for16S rRNA metabarcoding, **2.** Primers for the sequence determination of full-length 16S rRNA gene, **3.** Primers for PCR screening and sequencing of COTS27, **4.** Probes for FISH analysis.
