## Supplementary material for "A universal subcuticular bacterial symbiont of a coral predator, the crown-of-thorns starfish, in the Indo-Pacific": Suppl. table S1

### Supplementary Table S1

#### A universal subcuticular bacterial symbiont of a coral predator, the crown-of-thorns starfish, in the Indo-Pacific

Naohisa WADA, Hideaki YUASA, Rei KAJITANI, Yasuhiro GOTOH, Yoshitoshi OGURA, Dai YOSHIMURA, Atsushi TOYODA, Sen-Lin TANG, Yukio HIGASHIMURA, Hugh SWEATMAN, Zac FORSMAN, Omri BRONSTEIN, Gal EYAL, Nalinee THONGTHAM, Takehiko ITOH, Tetsuya HAYASHI, Nina YASUDA

**Suppl. Table S1** Sample number of body parts from COTS in the 16S rRNA metabarcoding analysis (total 130 samples)

| | Okinawa | | | Miyazaki | | |
|---|---|---|---|---|---|---|
| | **Okinawa1** | **Okinawa2** | **Okinawa3** | **Miyazaki1** | **Miyazaki2** | **Miyazaki3** |
| **Surface body parts** | | | | | | |
| **Aboral side** | | | | | | |
| **Disc spines** | | | | | | |
| **Tips** | 3 | 3 | 3 | 3 | 3 | 3 |
| **Bases** | 3 | 3 | 3 | 2 | 3 | 3 |
| **Arm spines** | | | | | | |
| **Tips** | 3 | 3 | 3 | 3 | 2 | 3 |
| **Bases** | 3 | 3 | 3 | 3 | 3 | 3 |
| **Oral side** | | | | | | |
| Ambulacral spines | | | | | | |
| **Tips** | 3 | 3 | 3 | – | – | – |
| **Bases** | 3 | 3 | 3 | – | – | – |
| **Whole** | – | – | – | 2 | 2 | 2 |
| **Tube foot** | 3 | 3 | 3 | 3 | 3 | 3 |
| **Internal body parts** | | | | | | |
| **Pyloric stomachs** | 3 | 3 | 3 | 3 | 3 | 3 |
| **Total** | 24 | 24 | 24 | 19 | 19 | 20 |
