## Supplementary material for "A universal subcuticular bacterial symbiont of a coral predator, the crown-of-thorns starfish, in the Indo-Pacific": Suppl. table S2

### Supplementary Table 2

#### A universal subcuticular bacterial symbiont of a coral predator, the crown-of-thorns starfish, in the Indo-Pacific

Naohisa WADA, Hideaki YUASA, Rei KAJITANI, Yasuhiro GOTOH, Yoshitoshi OGURA, Dai YOSHIMURA, Atsushi TOYODA, Sen-Lin TANG, Yukio HIGASHIMURA, Hugh SWEATMAN, Zac FORSMAN, Omri BRONSTEIN, Gal EYAL, Nalinee THONGTHAM, Takehiko ITOH, Tetsuya HAYASHI, Nina YASUDA

**Supp. table S2** Number of OTUs and sequence reads obtained from the 16S rRNA metabarcoding

| Sample ID | Seq. ID | Total | | Archaea | | Eukaryota | | Chloroplast | | Unknown | | Bacteria | | |
|---|---|---|---|---|---|---|---|---|---|---|---|---|---|---|
| | | OTUs | Reads | OTUs | Reads | OTUs | Reads | OTUs | Reads | OTUs | Reads | OTUs | Reads | OTU1 |
| Okinawa1_Disk Spine tip 1 | O1Dt1 | 67 | 17783 | 0 | 0 | 37 | 1158 | 0 | 0 | 4 | 1951 | 26 | 14674 | 10217 |
| Okinawa1_Disk Spine tip 2 | O1Dt2 | 146 | 15633 | 0 | 0 | 18 | 310 | 8 | 250 | 6 | 425 | 114 | 14648 | 2354 |
| Okinawa1_Disk Spine tip 3 | O1Dt3 | 72 | 21542 | 0 | 0 | 23 | 1018 | 0 | 0 | 7 | 1584 | 42 | 18940 | 7265 |
| Okinawa2_Disk Spine tip 1 | O2Dt1 | 19 | 23861 | 0 | 0 | 0 | 0 | 0 | 0 | 0 | 0 | 19 | 23861 | 0 |
| Okinawa2_Disk Spine tip 2 | O2Dt2 | 94 | 11125 | 0 | 0 | 18 | 39 | 5 | 297 | 4 | 563 | 67 | 10226 | 4723 |
| Okinawa2_Disk Spine tip 3 | O2Dt3 | 63 | 16127 | 0 | 0 | 32 | 106 | 0 | 0 | 4 | 1609 | 27 | 14412 | 12027 |
| Okinawa3_Disk Spine tip 1 | O3Dt1 | 84 | 11569 | 0 | 0 | 18 | 51 | 1 | 10 | 4 | 667 | 61 | 10841 | 7154 |
| Okinawa3_Disk Spine tip 2 | O3Dt2 | 114 | 13252 | 0 | 0 | 16 | 36 | 4 | 26 | 4 | 708 | 90 | 12482 | 3301 |
| Okinawa3_Disk Spine tip 3 | O3Dt3 | 72 | 16091 | 0 | 0 | 14 | 63 | 0 | 0 | 5 | 2564 | 53 | 13464 | 10000 |
| Okinawa1_Disk Spine base 1 | O1Db1 | 89 | 17899 | 0 | 0 | 53 | 2237 | 0 | 0 | 7 | 3184 | 29 | 12478 | 11470 |
| Okinawa1_Disk Spine base 2 | O1Db2 | 85 | 17065 | 0 | 0 | 53 | 2132 | 0 | 0 | 4 | 2802 | 28 | 12131 | 10835 |
| Okinawa1_Disk Spine base 3 | O1Db3 | 83 | 17203 | 0 | 0 | 52 | 2115 | 0 | 0 | 7 | 3173 | 24 | 11915 | 10734 |
| Okinawa2_Disk Spine base 1 | O2Db1 | 32 | 8870 | 0 | 0 | 8 | 20 | 0 | 0 | 3 | 628 | 21 | 8222 | 6920 |
| Okinawa2_Disk Spine base 2 | O2Db2 | 37 | 15995 | 0 | 0 | 10 | 23 | 0 | 0 | 3 | 1212 | 24 | 14760 | 11031 |
| Okinawa2_Disk Spine base 3 | O2Db3 | 17 | 17591 | 0 | 0 | 1 | 376 | 0 | 0 | 0 | 0 | 16 | 17215 | 0 |
| Okinawa3_Disk Spine base 1 | O3Db1 | 37 | 22515 | 0 | 0 | 3 | 12 | 0 | 0 | 4 | 5729 | 30 | 16774 | 10924 |
| Okinawa3_Disk Spine base 2 | O3Db2 | 47 | 23933 | 0 | 0 | 12 | 36 | 0 | 0 | 4 | 3252 | 31 | 20645 | 16772 |
| Okinawa3_Disk Spine base 3 | O3Db3 | 65 | 15928 | 0 | 0 | 26 | 150 | 0 | 0 | 7 | 1461 | 32 | 14317 | 11485 |
| Miyazaki1_Disk Spine tip 1 | M1Dt1 | 69 | 16664 | 0 | 0 | 31 | 196 | 0 | 0 | 5 | 5783 | 33 | 10685 | 6217 |

| | | | | | | | | | | | | | | |
|---|---|---|---|---|---|---|---|---|---|---|---|---|---|---|
| Miyazaki1_Disk Spine tip 2 | M1Dt2 | 79 | 11535 | 0 | 0 | 36 | 148 | 0 | 0 | 9 | 5122 | 34 | 6265 | 4734 |
| Miyazaki1_Disk Spine tip 3 | M1Dt3 | 90 | 18574 | 0 | 0 | 26 | 169 | 16 | 786 | 6 | 9149 | 42 | 8470 | 3769 |
| Miyazaki2_Disk Spine tip 1 | M2Dt1 | 111 | 14465 | 0 | 0 | 13 | 31 | 16 | 1708 | 7 | 2621 | 75 | 10105 | 7699 |
| Miyazaki2_Disk Spine tip 2 | M2Dt2 | 63 | 14781 | 2 | 20 | 17 | 29 | 2 | 19 | 3 | 1991 | 39 | 12722 | 11338 |
| Miyazaki2_Disk Spine tip 3 | M2Dt3 | 54 | 16897 | 1 | 2 | 1 | 4 | 2 | 62 | 3 | 2058 | 47 | 14771 | 12064 |
| Miyazaki3_Disk Spine tip 1 | M3Dt1 | 42 | 14456 | 0 | 0 | 15 | 28 | 0 | 0 | 3 | 1478 | 24 | 12950 | 11823 |
| Miyazaki3_Disk Spine tip 2 | M3Dt2 | 57 | 15567 | 0 | 0 | 16 | 24 | 3 | 10 | 3 | 1229 | 35 | 14304 | 13683 |
| Miyazaki3_Disk Spine tip 3 | M3Dt3 | 76 | 18549 | 0 | 0 | 24 | 108 | 6 | 50 | 6 | 2931 | 40 | 15460 | 14108 |
| Miyazaki1_Disk Spine base 2 | M1Db2 | 33 | 17864 | 0 | 0 | 2 | 28 | 0 | 0 | 3 | 5155 | 28 | 12681 | 5003 |
| Miyazaki1_Disk Spine base 3 | M1Db3 | 34 | 23321 | 0 | 0 | 1 | 10 | 2 | 80 | 3 | 5872 | 28 | 17359 | 7552 |
| Miyazaki2_Disk Spine base 1 | M2Db1 | 38 | 18732 | 1 | 5 | 1 | 3 | 2 | 10 | 3 | 2478 | 31 | 16236 | 11125 |
| Miyazaki2_Disk Spine base 2 | M2Db2 | 42 | 12599 | 0 | 0 | 8 | 15 | 0 | 0 | 3 | 2646 | 31 | 9938 | 8823 |
| Miyazaki2_Disk Spine base 3 | M2Db3 | 38 | 16239 | 0 | 0 | 2 | 3 | 1 | 1 | 3 | 1800 | 32 | 14435 | 10015 |
| Miyazaki3_Disk Spine base 1 | M3Db1 | 44 | 15517 | 0 | 0 | 2 | 8 | 4 | 43 | 3 | 1109 | 35 | 14357 | 10086 |
| Miyazaki3_Disk Spine base 2 | M3Db2 | 38 | 15143 | 0 | 0 | 7 | 24 | 0 | 0 | 3 | 1491 | 28 | 13628 | 10855 |
| Miyazaki3_Disk Spine base 3 | M3Db3 | 34 | 16885 | 0 | 0 | 7 | 30 | 0 | 0 | 4 | 1740 | 23 | 15115 | 12990 |
| Okinawa1_Arm Spine tip 1 | O1At1 | 60 | 17906 | 0 | 0 | 32 | 1336 | 0 | 0 | 3 | 2093 | 25 | 14477 | 12028 |
| Okinawa1_Arm Spine tip 2 | O1At2 | 56 | 13628 | 0 | 0 | 38 | 1436 | 0 | 0 | 5 | 2221 | 13 | 9971 | 9203 |
| Okinawa1_Arm Spine tip 3 | O1At3 | 44 | 17483 | 0 | 0 | 24 | 1199 | 0 | 0 | 2 | 1853 | 18 | 14431 | 8699 |
| Okinawa2_Arm Spine tip 1 | O2At1 | 101 | 16407 | 0 | 0 | 32 | 125 | 3 | 49 | 4 | 2343 | 62 | 13890 | 6023 |
| Okinawa2_Arm Spine tip 2 | O2At2 | 63 | 12766 | 0 | 0 | 32 | 100 | 0 | 0 | 5 | 1801 | 26 | 10865 | 9339 |
| Okinawa2_Arm Spine tip 3 | O2At3 | 68 | 20942 | 0 | 0 | 34 | 173 | 0 | 0 | 6 | 3823 | 28 | 16946 | 14171 |

| | | | | | | | | | | | | | | |
|---|---|---|---|---|---|---|---|---|---|---|---|---|---|---|
| Okinawa3_Arm Spine tip 1 | O3At1 | 78 | 14624 | 0 | 0 | 26 | 98 | 0 | 0 | 5 | 2367 | 47 | 12159 | 10185 |
| Okinawa3_Arm Spine tip 2 | O3At2 | 25 | 11307 | 0 | 0 | 2 | 70 | 0 | 0 | 4 | 24 | 19 | 11213 | 0 |
| Okinawa3_Arm Spine tip 3 | O3At3 | 56 | 17211 | 0 | 0 | 20 | 95 | 0 | 0 | 5 | 2964 | 31 | 14152 | 11949 |
| Okinawa1_Arm Spine base 1 | O1Ab1 | 60 | 16033 | 0 | 0 | 33 | 1533 | 0 | 0 | 3 | 2239 | 24 | 12261 | 10933 |
| Okinawa1_Arm Spine base 2 | O1Ab2 | 76 | 16995 | 0 | 0 | 48 | 2350 | 0 | 0 | 8 | 3591 | 20 | 11054 | 10025 |
| Okinawa1_Arm Spine base 3 | O1Ab3 | 53 | 17307 | 0 | 0 | 30 | 911 | 0 | 0 | 5 | 1233 | 18 | 15163 | 14136 |
| Okinawa2_Arm Spine base 1 | O2Ab1 | 65 | 16333 | 0 | 0 | 30 | 120 | 0 | 0 | 3 | 1404 | 32 | 14809 | 11674 |
| Okinawa2_Arm Spine base 2 | O2Ab2 | 55 | 19710 | 0 | 0 | 15 | 45 | 0 | 0 | 6 | 1119 | 34 | 18546 | 15875 |
| Okinawa2_Arm Spine base 3 | O2Ab3 | 45 | 17408 | 0 | 0 | 18 | 32 | 0 | 0 | 4 | 1157 | 23 | 16219 | 5834 |
| Okinawa3_Arm Spine base 1 | O3Ab1 | 45 | 12844 | 0 | 0 | 6 | 12 | 0 | 0 | 5 | 1430 | 34 | 11402 | 6866 |
| Okinawa3_Arm Spine base 2 | O3Ab2 | 30 | 14578 | 0 | 0 | 5 | 9 | 0 | 0 | 4 | 2044 | 21 | 12525 | 8809 |
| Okinawa3_Arm Spine base 3 | O3Ab3 | 31 | 14994 | 0 | 0 | 7 | 88 | 0 | 0 | 4 | 2321 | 20 | 12585 | 9798 |
| Miyazaki1_Arm Spine tip 1 | M1At1 | 88 | 15614 | 1 | 31 | 42 | 235 | 5 | 74 | 6 | 4567 | 34 | 10707 | 6501 |
| Miyazaki1_Arm Spine tip 2 | M1At2 | 92 | 14205 | 0 | 0 | 48 | 295 | 2 | 14 | 9 | 6238 | 33 | 7658 | 5298 |
| Miyazaki1_Arm Spine tip 3 | M1At3 | 54 | 18626 | 0 | 0 | 17 | 111 | 2 | 110 | 6 | 5163 | 29 | 13242 | 10516 |
| Miyazaki2_Arm Spine tip 1 | M2At1 | 118 | 15769 | 7 | 80 | 1 | 2 | 18 | 823 | 4 | 1689 | 88 | 13175 | 9326 |
| Miyazaki2_Arm Spine tip 3 | M2At3 | 57 | 16220 | 0 | 0 | 23 | 53 | 0 | 0 | 7 | 2665 | 27 | 13502 | 12932 |
| Miyazaki3_Arm Spine tip 1 | M3At1 | 66 | 16590 | 0 | 0 | 26 | 72 | 2 | 3 | 4 | 1599 | 34 | 14916 | 11060 |
| Miyazaki3_Arm Spine tip 2 | M3At2 | 73 | 13477 | 0 | 0 | 35 | 85 | 4 | 10 | 4 | 1888 | 30 | 11494 | 8937 |
| Miyazaki3_Arm Spine tip 3 | M3At3 | 60 | 16339 | 0 | 0 | 23 | 72 | 0 | 0 | 6 | 2459 | 31 | 13808 | 11239 |
| Miyazaki1_Arm Spine base 1 | M1Ab1 | 35 | 16398 | 0 | 0 | 5 | 42 | 0 | 0 | 4 | 5281 | 26 | 11075 | 5266 |
| Miyazaki1_Arm Spine base 2 | M1Ab2 | 44 | 14705 | 0 | 0 | 5 | 51 | 3 | 12 | 3 | 3013 | 33 | 11629 | 7010 |

| | | | | | | | | | | | | | | |
|---|---|---|---|---|---|---|---|---|---|---|---|---|---|---|
| Miyazaki1_Arm Spine base 3 | M1Ab3 | 31 | 18591 | 0 | 0 | 3 | 7 | 0 | 0 | 4 | 3071 | 24 | 15513 | 3604 |
| Miyazaki2_Arm Spine base 1 | M2Ab1 | 36 | 14550 | 0 | 0 | 2 | 6 | 0 | 0 | 3 | 3421 | 31 | 11123 | 9826 |
| Miyazaki2_Arm Spine base 2 | M2Ab2 | 27 | 12266 | 0 | 0 | 1 | 1 | 1 | 5 | 3 | 2484 | 22 | 9776 | 8994 |
| Miyazaki2_Arm Spine base 3 | M2Ab3 | 39 | 14460 | 1 | 2 | 1 | 1 | 4 | 8 | 3 | 1675 | 30 | 12774 | 11399 |
| Miyazaki3_Arm Spine base 1 | M3Ab1 | 21 | 15976 | 0 | 0 | 0 | 0 | 0 | 0 | 1 | 1 | 20 | 15975 | 1 |
| Miyazaki3_Arm Spine base 2 | M3Ab2 | 35 | 19258 | 0 | 0 | 1 | 55 | 0 | 0 | 3 | 10198 | 31 | 9005 | 123 |
| Miyazaki3_Arm Spine base 3 | M3Ab3 | 72 | 19177 | 2 | 16 | 5 | 29 | 3 | 34 | 3 | 2295 | 59 | 16803 | 11694 |
| Okinawa1_Ambulacral Spine tip1 | O1Bt1 | 83 | 17618 | 1 | 40 | 29 | 571 | 4 | 155 | 5 | 667 | 44 | 16185 | 10721 |
| Okinawa1_Ambulacral Spine tip2 | O1Bt2 | 47 | 12528 | 0 | 0 | 19 | 852 | 0 | 0 | 5 | 1218 | 23 | 10458 | 7529 |
| Okinawa1_Ambulacral Spine tip3 | O1Bt3 | 72 | 17616 | 0 | 0 | 37 | 2210 | 3 | 109 | 8 | 3097 | 24 | 12200 | 8313 |
| Okinawa2_Ambulacral Spine tip 1 | O2Bt1 | 54 | 15746 | 0 | 0 | 21 | 88 | 0 | 0 | 5 | 2470 | 28 | 13188 | 6848 |
| Okinawa2_Ambulacral Spine tip 2 | O2Bt2 | 92 | 12408 | 0 | 0 | 18 | 64 | 7 | 68 | 4 | 715 | 63 | 11561 | 6302 |
| Okinawa2_Ambulacral Spine tip 3 | O2Bt3 | 61 | 16696 | 0 | 0 | 29 | 120 | 0 | 0 | 7 | 2262 | 25 | 14314 | 7911 |
| Okinawa3_Ambulacral Spine tip 1 | O3Bt1 | 115 | 12372 | 0 | 0 | 42 | 236 | 1 | 6 | 7 | 3458 | 65 | 8672 | 5184 |
| Okinawa3_Ambulacral Spine tip 2 | O3Bt2 | 72 | 9325 | 0 | 0 | 37 | 124 | 1 | 2 | 5 | 1703 | 29 | 7496 | 6475 |
| Okinawa3_Ambulacral Spine tip 3 | O3Bt3 | 100 | 13316 | 0 | 0 | 43 | 241 | 0 | 0 | 7 | 2857 | 50 | 10218 | 8401 |
| Okinawa1_Ambulacral Spine base 1 | O1Bb1 | 62 | 15033 | 0 | 0 | 44 | 969 | 0 | 0 | 2 | 1273 | 16 | 12791 | 11322 |
| Okinawa1_Ambulacral Spine base 2 | O1Bb2 | 58 | 17913 | 0 | 0 | 32 | 1112 | 0 | 0 | 2 | 1562 | 24 | 15239 | 11836 |
| Okinawa1_Ambulacral Spine base 3 | O1Bb3 | 67 | 16846 | 0 | 0 | 37 | 1058 | 0 | 0 | 3 | 1400 | 27 | 14388 | 11108 |
| Okinawa2_Ambulacral Spine base 1 | O2Bb1 | 61 | 18228 | 0 | 0 | 28 | 115 | 2 | 10 | 4 | 1167 | 27 | 16936 | 11036 |
| Okinawa2_Ambulacral Spine base 2 | O2Bb2 | 51 | 19758 | 0 | 0 | 17 | 51 | 0 | 0 | 4 | 1746 | 30 | 17961 | 12001 |
| Okinawa2_Ambulacral Spine base 3 | O2Bb3 | 53 | 21947 | 0 | 0 | 23 | 118 | 0 | 0 | 4 | 2267 | 26 | 19562 | 11438 |

| | | | | | | | | | | | | | | |
|---|---|---|---|---|---|---|---|---|---|---|---|---|---|---|
| Okinawa3_Ambulacral Spine base 1 | O3Bb1 | 54 | 2592 | 0 | 0 | 24 | 79 | 0 | 0 | 5 | 1019 | 25 | 1494 | 579 |
| Okinawa3_Ambulacral Spine base 2 | O3Bb2 | 58 | 11467 | 0 | 0 | 34 | 121 | 0 | 0 | 3 | 1929 | 21 | 9417 | 7789 |
| Okinawa3_Ambulacral Spine base 3 | O3Bb3 | 74 | 11681 | 0 | 0 | 37 | 183 | 0 | 0 | 7 | 3247 | 30 | 8251 | 5404 |
| Miyazaki1_Ambulacral Spine 1 | M1Bw1 | 41 | 15418 | 1 | 13 | 3 | 9 | 1 | 1 | 3 | 2879 | 33 | 12516 | 11871 |
| Miyazaki1_Ambulacral Spine 2 | M1Bw2 | 59 | 14110 | 0 | 0 | 25 | 125 | 1 | 11 | 7 | 6033 | 26 | 7941 | 7450 |
| Miyazaki2_Ambulacral Spine 1 | M2Bw1 | 46 | 13053 | 0 | 0 | 14 | 19 | 0 | 0 | 3 | 2670 | 29 | 10364 | 9950 |
| Miyazaki2_Ambulacral Spine 2 | M2Bw2 | 62 | 14415 | 1 | 7 | 32 | 85 | 1 | 2 | 4 | 3123 | 24 | 11198 | 10729 |
| Miyazaki3_Ambulacral Spine 1 | M3Bw1 | 79 | 11965 | 1 | 16 | 30 | 93 | 5 | 30 | 4 | 2384 | 39 | 9442 | 7657 |
| Miyazaki3_Ambulacral Spine 2 | M3Bw2 | 71 | 15123 | 0 | 0 | 35 | 98 | 0 | 0 | 6 | 3209 | 30 | 11816 | 10272 |
| Okinawa1_Tube feet 1 | O1T1 | 75 | 9470 | 0 | 0 | 54 | 1701 | 0 | 0 | 4 | 2392 | 17 | 5377 | 5175 |
| Okinawa1_Tube feet 2 | O1T2 | 74 | 9160 | 0 | 0 | 49 | 1403 | 0 | 0 | 4 | 2045 | 21 | 5712 | 4693 |
| Okinawa1_Tube feet 3 | O1T3 | 65 | 11697 | 0 | 0 | 42 | 991 | 0 | 0 | 3 | 1362 | 20 | 9344 | 9044 |
| Okinawa2_Tube feet 1 | O2T1 | 82 | 14484 | 0 | 0 | 43 | 188 | 0 | 0 | 9 | 2066 | 30 | 12230 | 9646 |
| Okinawa2_Tube feet 2 | O2T2 | 69 | 16941 | 0 | 0 | 15 | 35 | 0 | 0 | 5 | 243 | 49 | 16663 | 1362 |
| Okinawa2_Tube feet 3 | O2T3 | 96 | 13831 | 0 | 0 | 59 | 467 | 0 | 0 | 10 | 4214 | 27 | 9150 | 6024 |
| Okinawa3_Tube feet 1 | O3T1 | 97 | 12706 | 0 | 0 | 45 | 320 | 0 | 0 | 6 | 5404 | 46 | 6982 | 4753 |
| Okinawa3_Tube feet 2 | O3T2 | 78 | 9280 | 0 | 0 | 47 | 345 | 0 | 0 | 7 | 2498 | 24 | 6437 | 5609 |
| Okinawa3_Tube feet 3 | O3T3 | 99 | 10321 | 0 | 0 | 65 | 592 | 1 | 20 | 8 | 4540 | 25 | 5169 | 3992 |
| Miyazaki1_Tube feet 1 | M1T1 | 71 | 15623 | 0 | 0 | 27 | 130 | 0 | 0 | 7 | 4952 | 37 | 10541 | 9057 |
| Miyazaki1_Tube feet 2 | M1T2 | 56 | 15101 | 0 | 0 | 19 | 72 | 1 | 1 | 6 | 3345 | 30 | 11683 | 9770 |
| Miyazaki1_Tube feet 3 | M1T3 | 45 | 16813 | 0 | 0 | 10 | 69 | 0 | 0 | 4 | 3418 | 31 | 13326 | 10008 |
| Miyazaki2_Tube feet 1 | M2T1 | 109 | 16685 | 0 | 0 | 62 | 642 | 0 | 0 | 13 | 10012 | 34 | 6031 | 5146 |

| | | | | | | | | | | | | | | |
|---|---|---|---|---|---|---|---|---|---|---|---|---|---|---|
| Miyazaki2_Tube feet 2 | M2T2 | 91 | 7365 | 0 | 0 | 58 | 331 | 0 | 0 | 8 | 4434 | 25 | 2600 | 2391 |
| Miyazaki2_Tube feet 3 | M2T3 | 100 | 13669 | 0 | 0 | 63 | 369 | 0 | 0 | 10 | 6649 | 27 | 6651 | 5977 |
| Miyazaki3_Tube feet 1 | M3T1 | 71 | 11791 | 0 | 0 | 41 | 108 | 0 | 0 | 4 | 2136 | 26 | 9547 | 8597 |
| Miyazaki3_Tube feet 2 | M3T2 | 59 | 12925 | 0 | 0 | 23 | 62 | 0 | 0 | 4 | 1826 | 32 | 11037 | 9799 |
| Miyazaki3_Tube feet 3 | M3T3 | 47 | 14942 | 0 | 0 | 12 | 20 | 0 | 0 | 3 | 2415 | 32 | 12507 | 9043 |
| Okinawa1_Stomach 1 | O1S1 | 96 | 7968 | 0 | 0 | 69 | 3146 | 0 | 0 | 6 | 4099 | 21 | 723 | 12 |
| Okinawa1_Stomach 2 | O1S2 | 105 | 7511 | 0 | 0 | 71 | 3031 | 0 | 0 | 9 | 3777 | 25 | 703 | 1 |
| Okinawa1_Stomach 3 | O1S3 | 37 | 16731 | 0 | 0 | 9 | 1270 | 0 | 0 | 3 | 1903 | 25 | 13558 | 1 |
| Okinawa2_Stomach 1 | O2S1 | 107 | 7061 | 0 | 0 | 75 | 688 | 0 | 0 | 10 | 5860 | 22 | 513 | 59 |
| Okinawa2_Stomach 2 | O2S2 | 92 | 7927 | 0 | 0 | 62 | 597 | 0 | 0 | 7 | 5162 | 23 | 2168 | 1306 |
| Okinawa2_Stomach 3 | O2S3 | 108 | 11561 | 0 | 0 | 78 | 788 | 0 | 0 | 8 | 9540 | 22 | 1233 | 146 |
| Okinawa3_Stomach 1 | O3S1 | 100 | 9142 | 0 | 0 | 73 | 795 | 0 | 0 | 9 | 5506 | 18 | 2841 | 1 |
| Okinawa3_Stomach 2 | O3S2 | 97 | 10874 | 0 | 0 | 70 | 527 | 0 | 0 | 8 | 3841 | 19 | 6506 | 1 |
| Okinawa3_Stomach 3 | O3S3 | 103 | 9945 | 0 | 0 | 73 | 766 | 1 | 10 | 7 | 7519 | 22 | 1650 | 10 |
| Miyazaki1_Stomach 1 | M1S1 | 114 | 15446 | 0 | 0 | 60 | 843 | 1 | 1 | 12 | 13128 | 41 | 1474 | 148 |
| Miyazaki1_Stomach 2 | M1S2 | 139 | 14178 | 4 | 22 | 53 | 738 | 15 | 125 | 8 | 12258 | 59 | 1035 | 53 |
| Miyazaki1_Stomach 3 | M1S3 | 104 | 19969 | 0 | 0 | 51 | 447 | 0 | 0 | 9 | 15890 | 44 | 3632 | 30 |
| Miyazaki2_Stomach 1 | M2S1 | 107 | 14303 | 0 | 0 | 72 | 686 | 0 | 0 | 10 | 12573 | 25 | 1044 | 73 |
| Miyazaki2_Stomach 2 | M2S2 | 140 | 15132 | 0 | 0 | 81 | 829 | 4 | 10 | 12 | 13677 | 43 | 616 | 25 |
| Miyazaki2_Stomach 3 | M2S3 | 129 | 20375 | 1 | 7 | 64 | 570 | 6 | 67 | 8 | 18547 | 50 | 1184 | 27 |
| Miyazaki3_Stomach 1 | M3S1 | 99 | 14411 | 2 | 3 | 32 | 526 | 2 | 3 | 6 | 11820 | 57 | 2059 | 276 |
| Miyazaki3_Stomach 2 | M3S2 | 65 | 14795 | 1 | 2 | 19 | 448 | 1 | 1 | 5 | 11626 | 39 | 2718 | 353 |

| | | | | | | | | | | | | | | |
|---|---|---|---|---|---|---|---|---|---|---|---|---|---|---|
| Miyazaki3_Stomach 3 | M3S3 | 109 | 25351 | 0 | 0 | 43 | 507 | 0 | 0 | 6 | 21855 | 60 | 2989 | 77 |
| Seawater_Okinawa1 | OSW1 | 255 | 18533 | 12 | 667 | 0 | 0 | 29 | 1187 | 1 | 1 | 213 | 16678 | 5 |
| Seawater_Okinawa2 | OSW2 | 268 | 23260 | 10 | 954 | 0 | 0 | 29 | 1446 | 1 | 1 | 228 | 20859 | 6 |
| Seawater_Okinawa3 | OSW3 | 282 | 23807 | 12 | 674 | 0 | 0 | 30 | 1473 | 1 | 1 | 239 | 21659 | 4 |
| Seawater_Miyazaki1 | MSW1 | 343 | 19960 | 26 | 2551 | 0 | 0 | 40 | 2218 | 1 | 1 | 276 | 15190 | 0 |
| Seawater_Miyazaki2 | MSW2 | 355 | 21131 | 26 | 3370 | 0 | 0 | 43 | 3074 | 2 | 3 | 284 | 14684 | 0 |
| Seawater_Miyazaki3 | MSW3 | 352 | 28661 | 26 | 4204 | 0 | 0 | 43 | 5191 | 1 | 2 | 282 | 19264 | 0 |
