## Supplementary material for "A universal subcuticular bacterial symbiont of a coral predator, the crown-of-thorns starfish, in the Indo-Pacific": Suppl. table S3-9

### Supplementary Tables S3-9

#### A universal subcuticular bacterial symbiont of a coral predator, the crown-of-thorns starfish, in the Indo-Pacific

Naohisa WADA, Hideaki YUASA, Rei KAJITANI, Yasuhiro GOTOH, Yoshitoshi OGURA, Dai YOSHIMURA, Atsushi TOYODA, Sen-Lin TANG, Yukio HIGASHIMURA, Hugh SWEATMAN, Zac FORSMAN, Omri BRONSTEIN, Gal EYAL, Nalinee THONGTHAM, Takehiko ITOH, Tetsuya HAYASHI, Nina YASUDA

**Suppl. table S3 Bacterial taxa number identified from phylum to family in the COTSs and seawater samples (total 761 OTUs).**

| | COTS | | | Seawater | | |
|---|---|---|---|---|---|---|
| | Total | Okinawan individuals (n=3) | Miyazaki's individuals (n=3) | Total | Okinawan seawater (n=3) | Miyazaki's seawater (n=3) |
| Phylum | 19 | 15 | 17 | 16 | 15 | 15 |
| Class | 34 (1) | 26 | 28 (1) | 24 (1) | 21 (1) | 21 (1) |
| Order | 92 (6) | 68 (5) | 73 (4) | 68 (5) | 57 (5) | 62 (4) |
| Family | 144 (5) | 109 (1) | 110 (5) | 96 (3) | 79 (2) | 89 (3) |
| Unclassified phylum | 29 OTUs | 20 OTUs | 17 OTUs | 22 OTUs | 17 OTUs | 17 OTUs |
| Unknown kingdom [*1] | 7 OTUs | 3 OTUs | 5 OTUs | 12 OTUs | 11 OTUs | 10 OTUs |

**The number of bacterial taxa and their corresponding taxonomic level. The number of both unclassified or unknown bacterial taxa are mentioned in brackets.**

***1 The OTUs were classified as bacteria in the Silva SINA aligner (https://www.arb-silva.de/aligner/) after sorting in MOTHUR (https://mothur.org/).**

**Suppl. table S4** Abundance of COTS27 (OTU1) in the different body components of COTS among Okinawa and Miyazaki samples

| | **Average abundance of COTS27 (OTU1) (%)** | | |
|---|---|---|---|
| | **Total** | **Okinawa** | **Miyazaki** |
| **Surface body parts** | | | |
| **All spines** | **68.8 ± 24.1 (n=94)** | **66.5 ± 24.1 (n=54)** | **72.0 ± 24.1 (n=40)** |
| Aboral side | | | |
| Disc spines | 66.5 ± 26.0 (n=35) | 59.8 ± 30.9 (n=18) | 73.5 ± 17.8 (n=17) |
| Tips | 62.4 ± 28.1 (n=18) | 46.7 ± 28.7 (n=9) | 78.2 ± 17.0 (n=9) |
| Bases | 70.7 ± 23.7 (n=17) | 73.0 ± 28.6 (n=9) | 68.2 ± 18.2 (n=8) |
| Arm spines | 68.0 ± 26.9 (n=35) | 72.1 ± 24.4 (n=18) | 63.6 ± 29.5 (n=17) |
| Tips | 72.1 ± 22.6 (n=17) | 68.5 ± 30.0 (n=9) | 76.1 ± 10.3 (n=8) |
| Bases | 64.1 ± 30.6 (n=18) | 75.8 ± 18.3 (n=9) | 52.4 ± 36.7 (n=9) |
| Oral side | | | |
| Ambulacral spines | 73.6 ± 15.8 (n=24) | 67.6 ± 13.3 (n=18) | 91.4 ± 6.1 (n=6) |
| Tips | 66.3 ± 12.3 (n=9) | 66.3 ± 12.3 (n=9) | – |
| Bases | 69.0 ± 14.9 (n=9) | 69.0 ± 14.9 (n=9) | – |
| Whole | 91.4 ± 6.1 (n=6) | – | 91.4 ± 6.1 (n=6) |
| **Tube feet** | **79.1 ± 19.8 (n=18)** | **73.4 ± 26.7 (n=9)** | **84.8 ± 6.8 (n=9)** |
| **Internal body parts** | | | |
| **Pyloric stomachs** | **8.0 ± 13.9 (n=18)** | **9.6 ± 19.6 (n=9)** | **6.5 ± 4.7 (n=9)** |

**Suppl. table S5** COTS genome sequencing read data. All reads were preprocessed to exclude adaptor sequences and low-quality bases by Platanus_trim (version 1.0.7; http://platanus.bio.titech.ac.jp/). For all mate-pair libraries, short-insert pairs (estimated insert size ≤ 0.5 × nominal size) and PCR products, duplicates were removed based on mapping information to the assembled results (scaffolds), which were constructed only from the paired-end libraries, using an in-house program.

| | | Raw | | Pre-processed | |
|---|---|---|---|---|---|
| Library type | Nominal insert length (bp) | Mean read length (bp) | Total length (bp) | Mean read length (bp) | Total length (bp) |
| paired-ends | 300 | 150 | 33,596,575,800 | 147 | 32,610,796,353 |
| paired-ends | 500 | 150 | 34,403,911,800 | 147 | 32,812,079,166 |
| mate-pairs | 3,000 | 150 | 21,372,289,200 | 116 | 11,201,114,333 |
| mate-pairs | 5,000 | 150 | 22,326,121,800 | 118 | 11,998,456,502 |
| mate-pairs | 8,000 | 150 | 22,750,867,500 | 120 | 12,396,355,042 |
| mate-pairs | 10,000 | 150 | 22,390,767,000 | 121 | 11,476,430,100 |
| mate-pairs | 12,000 | 150 | 22,224,980,700 | 119 | 10,842,998,778 |
| mate-pairs | 15,000 | 150 | 24,490,445,400 | 120 | 11,070,146,690 |

**Suppl. table S6** COTS27 genome information

| COTS27 genome | |
|---|---|
| Genome size | 2,684,921 bp |
| Gaps | 23 |
| Total number of Ns | 392 bp |
| GC ratio | 39.59% |
| Proteins | 1,650 |
| tRNAs | 35 |
| rRNAs | 3 |

**Suppl. table S7** COTS27 genome general biosynthesis profile based on KEGG metabolic pathways

| Biosynthesis | States[*1] | Biosynthesis | States[*1] |
|---|---|---|---|
| ***Amino acid biosynthesis*** | | ***Metabolism of cofactors and vitamins*** | |
| Alanine | ++ | Thiamine | – |
| Arginine | ++ | Riboflavin [*2] | ++ |
| Asparagine | – | Pyridoxal | + |
| Aspartic acid | – | NAD | ++ |
| Cysteine | ++ | Pantothenate | – |
| Glutamine | ++ | Coenzyme A | ++ |
| Glutamic acid | ++ | Pimeloyl-ACP | – |
| Glycine | + | Biotin | ++ |
| Histidine | + | Tetrahydrofolate | + |
| Isoleucine | ++ | L-threo-Tetrahydrobiopterin | – |
| Leucine | ++ | C1-unit interconversion | ++ |
| Lysine | ++ | Hemo | – |
| Methionine | ++ | Siroheme | – |
| Phenylalanine | ++ | Menaquinone | – |
| Proline | ++ | ***Lipid metabolism*** | |
| Serine | + | ***•Fatty acid biosynthesis degradation*** | |
| Threonine | + | Fatty acid | ++ |
| Tryptophan | ++ | Beta-Oxidation | + |
| Tyrosine | ++ | ***•Lipid metabolism*** | |
| Valine | ++ | Phosphatidylethanolamine (PE) | ++ |
| ***Nucleotide biosynthesis*** | | Ketone body | – |
| ***•Purine*** | | Triacylglycerol | – |
| Inosine monophosphate | ++ | | |
| Adenine ribonucleotide | ++ | | |
| Guanine ribonucleotide | + | | |
| ***•Pyrimidine*** | | | |
| Uridine monophosphate | ++ | | |
| Pyrimidine ribonucleotide | ++ | | |

| | |
|---|---|
| Pyrimidine deoxy ribonucleotide | + + |

*1: Indication of the biosynthesis pathway prediction: complete (+ +), one block missing (+), and two or more blocks missing (–).

*2: Not including biosynthesis pathway of flavin adenine dinucleotide (FAD) and flavin mononucleotide (FMN).

**Suppl. table S8** Samples used for 16S rRNA metabarcoding, reconstruction of phylogenetic tree, and PCR screening and sequencing

| Analysis | Location I.D. | *n* [*1] | Location | Country | Year | Remarks |
|---|---|---|---|---|---|---|
| 1. 16S rRNA metabarcoding | | | | | | |
| | Miyazaki | 3 | Ohshima, Miyazaki | Japan | Nov. 2017 | This study |
| | Okinawa | 3 | Yamakawa, Okinawa | Japan | Jul. 2017 | This study |
| 2. Phylogenetic analysis of COTS27 | | | | | | |
| | Miyazaki | 3 | Ohshima, Miyazaki | Japan | Nov. 2017 | Same specimens with 1. |
| | Okinawa | 2 | Yamakawa, Okinawa | Japan | Jul. 2017 | Same specimens with 1. |
| 3. PCR screening and sequencing of COTS27 | | | | | | |
| | Wakayama | 16 | Wakayama | Japan | Dec. 2005–Jun. 2006 | [1] |
| | Tatsukushi | 6 | Kouchi | Japan | Nov. 2005 | [1] |
| | Sakura-jima | 9 | Kagoshima | Japan | 2004–2005 | [1] |
| | Amami-Ohshima | 6 | Kagoshima | Japan | May–Sep. 2004 | [1] |
| | Onna-village | 18 | Okinawa | Japan | Jun. 2013 | This study |
| | Kerama Island | 9 | Okinawa | Japan | Jul. 2004 | [2] |
| | Kume Island | 16 | Okinawa | Japan | Sep. 2005 | [1] |
| | Miyako Island | 6 | Okinawa | Japan | May–Sep. 2004 | [1] |
| | Sekisei Lagoon, | 12 | Okinawa | Japan | 2004 | This study |
| | Bowden Reef | 16 | Great Barrier Reef | Australia | Mar. 2007 | [1] |
| | Clack Reef | 18 | Great Barrier Reef | Australia | Nov. 2006 | [1] |
| | Shell Reef | 28 | Great Barrier Reef | Australia | Mar. 2007 | [1] |

| | | | | | | |
|---|---|---|---|---|---|---|
| | Hawaii | 23 | Hawaii | United States | 2007–Sep. 2014 | This study |
| | Phuket | 4 | Phuket | Thailand | Oct. 2017–Aug. 2017 | This study |
| | Eilat | 8 | Eilat | Israel | Feb. 2017 | This study |
| 4. Fluorescence *in situ* hybridization (FISH) | | | | | | |
| | Miyazaki | 3 | Ohshima, Miyazaki | Japan | Apr. 2017 | This study |
| 5. Hologenome sequencing analysis | | | | | | |
| | Miyazaki | 1 | Ohshima, Miyazaki | Japan | Aug. 2014 | This study |

*1 *n* = number of the individuals

References;

1. Yasuda N, Nagai S, Hamaguchi M, Okaji K, Gérard K, Nadaoka K. Gene flow of Acanthaster planci (L.) in relation to ocean currents revealed by microsatellite analysis. *Mol Ecol* 2009; **18**: 1574–1590.

2. Yasuda N, Ogasawara K, Kajiwara K, Ueno M, Oki K, Taniguchi H, et al. Latitudinal differentiation in the reproduction patterns of the crown-of-thorns starfish Acanthaster planci through the Ryukyu Island Archipelago. *Plankton Benthos Res* 2010; **5**: 156–164.

**Suppl. table S9** Primers used for 16S rRNA gene sequence analysis and probes for FISH used in this study

| Analysis | Name | Sequence (5 '–3 ') | Targeted organism | *E. coli* position | Ref. and remarks |
|---|---|---|---|---|---|
| 16S rRNA metabarcoding | 515F_EMP | TCGTCGGCAGCGTCAGATGTGTATAAGAGACAG-GTGYCAGCMGCCGCGGTAA | Most bacteria | 515 | [1, 2] |
| | 806rb_EMP | GTCTCGTGGGCTCGGAGATGTGTATAAGAGACAG-GGACTACNVGGGTWTCTAAT | Most bacteria | 806 | [1, 2] |
| Phylogenetic analysis of COTS27 using the full-length 16S rRNA gene sequence | 27F | AGAGTTTGATCMTGGCTCAG | Most bacteria | 27 | [3] |
| | COTS_V4_R | CCTACACCAGGAATTCCGACTA | COTS27 | 668 | This study |
| | COTS_V4R_F | TAGTCGGAATTCCTGGTGTAGG | COTS27 | 668 | This study |
| | 1492R(c) | TACGGTTACCTTGTTACGAC | Most bacteria | 1492 | [4] |
| | Microbiont F | GAGCAATCTCACATGGATGACG | COTS27 | 392 | This study |
| | Microbiont R | CATGCTGATCCGCGATTACTAG | COTS27 | 1342 | This study |
| PCR screening and sequencing of COTS27 | COTSsymb_F | GATAGCCGCTGTAATGGCGA | COTS27 | 148 | This study |
| | COTSsymb_R | AGGCCTTCGTCATCCATGTG | COTS27 | 399 | This study |
| | Hitode_16S_f*3 | TGACYGTGCRAAGGTAGRATAATCATTGC | *Asteroidea*-universal primer | – | This study |
| | Hitode_16S_r*3 | CGCTGTTATCCCTRYGGSAACTT | *Asteroidea*-universal | – | This study |

| | | | | | |
|---|---|---|---|---|---|
| | | | primer | | |
| FISH | COTSsymb*1 | CTCAGCGATGCTAACGCACC | COTS27 | 196 | This study, FA 30% *2 |
| | EUB338mix*1 | GCWGCCWCCCGTAGGWGT | Most bacteria | 338 | [5, 6], FA 30% *2 |
| | Non338[1] | ACATCCTACGGGAGGC | – | – | [7], FA 30% *2 |

*1. Probes included 5'-end fluorescein Cy3 labeled.

*2. Formamide (FA) concentration (v/v) for hybridization at 46˚C.

*3. Primers designed to amplify COTS mitochondrial 16S rRNA gene sequence as a positive control.

References;


1. Apprill A, McNally S, Parsons R, Weber L. Minor revision to V4 region SSU rRNA 806R gene primer greatly increases detection of SAR11 bacterioplankton. *Aquat Microb Ecol* 2015; **75**: 129–137.
2. Walters W, Hyde ER, Berg-Lyons D, Ackermann G, Humphrey G, Parada A, et al. Improved Bacterial 16S rRNA Gene (V4 and V4-5) and Fungal Internal Transcribed Spacer Marker Gene Primers for Microbial Community Surveys. *mSystems* 2016; **1**: e00009-15.
3. Lane D. 16S/23S rRNA sequencing. *Nucleic Acid Tech Bact Syst* 1991; 115–175.
4. Allen MA, Goh F, Burns BP, Neilan BA. Bacterial, archaeal and eukaryotic diversity of smooth and pustular microbial mat communities in the hypersaline lagoon of Shark Bay. *Geobiology* 2009; **7**: 82–96.
5. Daims H, Brühl A, Amann R, Schleifer KH, Wagner M. The domain-specific probe EUB338 is insufficient for the detection of all Bacteria: development and evaluation of a more comprehensive probe set. *Syst Appl Microbiol* 1999; **22**: 434–44.
6. Amann RI, Krumholz L, Stahl DA. Fluorescent-oligonucleotide probing of whole cells for determinative, phylogenetic, and

environmental studies in microbiology. *J Bacteriol* 1990; **172**: 762–770.

7. Wallner G, Amann R, Beisker W. Optimizing fluorescent in situ hybridization with rRNA-targeted oligonucleotide probes for flow cytometric identification of microorganisms. *Cytometry* 1993; **14**: 136–43.
